# Temperature Induced Codimension-One and Codimension-Two Bifurcations in Hodgkin-Huxley Neurons

**DOI:** 10.1101/2025.08.16.670668

**Authors:** Brandon Jungwoo Han, Arij Daou

**Affiliations:** Biomedical Engineering Program, American University of Beirut; School of Engineering, Stanford University; Department of Neuroscience: The University of Chicago

**Keywords:** Dynamical Systems Analysis, Bifurcation Analysis, Hopf Bifurcation, Hodgkin-Huxley Model, Neuronal Excitability, Temperature Dependence, Timescale Warping, Ion Channels

## Abstract

Temperature fluctuations can have detrimental effects on the firing pattern and electrical activity of biological neurons, eliciting diverse responses depending on the neuronal cell types and the underlying ion channels exhibited. Using the classical Hodgkin-Huxley (HH) model, we performed a comprehensive dynamical systems analysis to determine how temperature fluctuations alter neuronal excitability, spike morphology, and bifurcation structure. We first relied on experimentally-derived temperature coefficients, or Q10 values, associated with gating kinetics and conductances, and examined codimension-1 and codimension-2 bifurcations across a range of temperatures and standard HH parameters governing the intrinsic properties (firing frequency, spike amplitude, spike width, afterhyperpolarization (AHP), time-to-peak AHP, etc…) of the model HH neuron. Our analysis revealed that increasing temperature accelerates gating dynamics, leading to narrower and higher-frequency spikes but reduced amplitudes, and ultimately to a loss of sustained firing via temperature-induced depolarization block. We identified generalized Hopf (Bautin) bifurcations as critical boundaries beyond which the system becomes strictly monostable. Extending the model to independently scale sodium activation, sodium inactivation, and potassium activation kinetics showed that excitability is particularly sensitive to potassium gating dynamics. Our findings provide a quantitative framework for understanding temperature modulations of neuronal activity, highlighting how temperature reshapes the excitability landscape, unveiling the intricate interplays between the activation/inactivation kinetics of ion channels, and identifying key parameters governing temperature robustness in neuronal models.

## Introduction

Numerous studies have explored the impact of temperature on neuronal firing patterns across a range of animals, including squid, snails, lobsters, frogs, Aplysia, locusts, and crayfish [1–7]. These studies that examined the temperature dependence of intrinsic membrane properties and synaptic transmission in invertebrates [5,8,9], lower vertebrates [3,10] and mammalian neurons [11,12], have shown that temperature variations strongly influence biochemical reaction rates, affecting the dynamics of biological neurons where the embedded ion channels on their cell membranes are stochastic in nature. For example, cooling decreases the activation and deactivation rates of ion channels that govern neuron dynamics [13,14], slows down spike propagation along the axon [15–18], and reduces both synaptic and ion channel conductance [19,20].

Small physiological fluctuations in temperature can significantly alter neuronal physiology and function [21,22]. Studies conducted *in vitro* showed that warming hippocampal slices causes a transient increase in population spike amplitude [23,24], while higher temperatures (such as 40°C) cause the spreading of depression in the hippocampus [25,26]. Kim and Connors (2012) later observed that hyperthermia increases the excitability in both excitatory and inhibitory hippocampal neurons [12]. Moreover, cooling brain slices from 33°C to 27°C reversibly increases the input resistance, causes larger and wider action potentials, increases the afterhyperpolarization (AHP) amplitude, and increases spike-frequency adaptation [27]. Temperature increase has also been shown to induce higher firing rates in the pyloric rhythm [28,29] and mediates transitioning in neuronal firing modes from tonic to bursting [30] which can drastically change the information the neuron conveys to its network. Focal cooling of specific brain areas has also been used to establish causal links between brain regions and behaviors [29,31–39]. For example, cooling the premotor cortical-like nucleus (HVC) in the adult male zebra finch brain slows down the song tempo across all timescales by up to 45%, but only slightly alters the acoustic structure [33]. These finding necessitates the further explorations of temperature dependencies of the underlying neuronal processes.

In this work, we provide a theoretical framework for understanding how temperature influences the dynamics of Hodgkin-Huxley model neurons. First, the firing patterns and the intrinsic properties of model neurons under the different values of temperature are examined. Second, bifurcation diagrams and phase-plane analyses are discussed. Through the bifurcation analysis, we aimed to stratify the parameter spaces and identify the different attractors involved. Trajectories converging to point attractors (i.e. stable equilibria) represent neurons stabilizing at fixed membrane potentials, while stable periodic orbits are indicative of sustained chains of nerve impulses. By examining the transitions between these attractors and their associated trajectories, we are able to predict the role of temperature in modulating neuronal behavior, and assess the effects of temperature fluctuations on the various intrinsic properties of the model neuron. Finally, we discuss how our findings can be generalized to other models that incorporate temperature into the Hodgkin-Huxley model. The temperature sensitivity of a biological process is commonly quantified using a Q10 value, defined as the ratio of the rate of the process when the temperature changes by 10°C [15]. While most models account for temperature changes by adding this Q10 temperature-dependent exponential or linear multiplicative factor to the maximal conductance parameters (𝑔_𝑁𝑎_, 𝑔_𝐾_, and 𝑔_𝐿_) and/or gating variable kinetics (𝑚*′*, 𝑛*′*, and ℎ*′*), we were able to provide a more generalized understanding of how temperature influences the model’s dynamics by isolating the effects of these individual timescale distortions and analyzing the bifurcations they induce. Our work provides an elaborate dynamical systems analysis of the impact of temperature variations on spike frequency and the various intrinsic properties that HH model neurons exhibit.

## 2. Methods and Materials

We used the single-compartment conductance-based biophysical model developed by Hodgkin and Huxley in 1952 [40]. Model simulations and numerical analyses were conducted using XPPAUT (available at [https://sites.pitt.edu/∼phase/bard/bardware/xpp/xpp.html]) and the MATCONT package in MATLAB [41].

### 2.1 Computational Modeling

The HH model includes the spike-producing voltage-gated currents (IK and INa) as well as a leak current (IL). The functional forms of activation/inactivation functions, time constants as well as the differential equations used are illustrated below. The membrane potential of the model neuron satisfies the current balance equation:

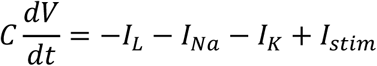

where 𝐼_𝑠𝑡𝑖𝑚_ represents the applied current and 𝐶 denotes membrane capacitance.

The leak current has a constant-conductance, while the voltage-dependent currents have non-constant conductances with activation/inactivation kinetics governed by voltage-dependent gating variables as described below. The voltage-dependent currents are:

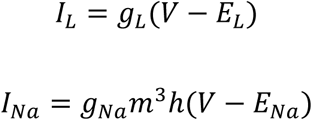

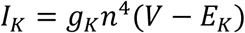

Here, each of the gating variable 𝑚, 𝑛, and ℎ takes values between 0 and 1 and satisfies a first-order differential equation of the form:

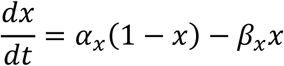

where 𝑥 = 𝑚, 𝑛, or ℎ.

At a constant membrane potential, the gating variables converge to a steady state, expressed as:

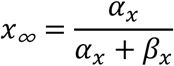

with a time constant given by:

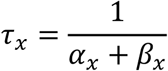

The model parameters as well as the functional forms for each of the activation variables are shown in Tables 1 and 2, respectively.

**Table 1.**
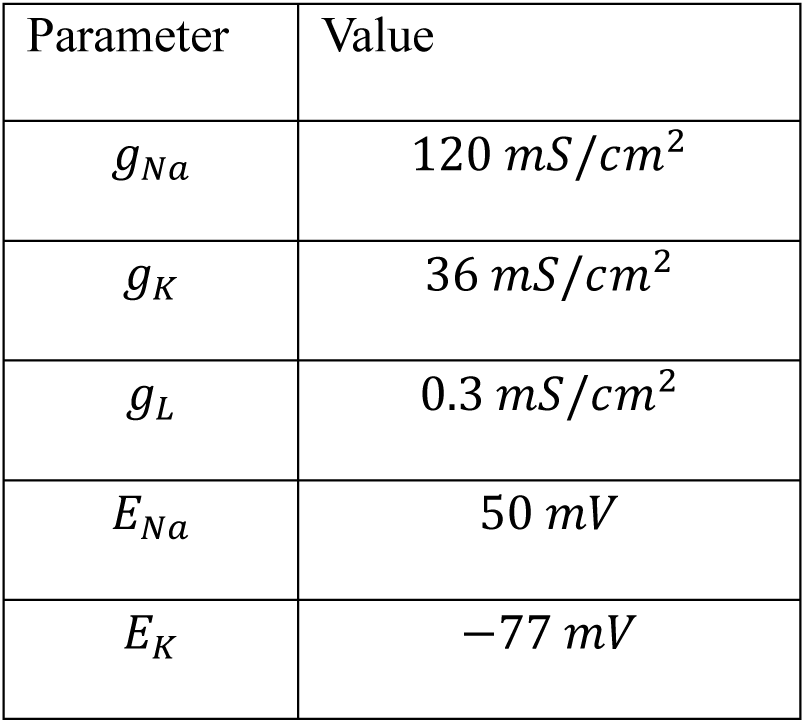
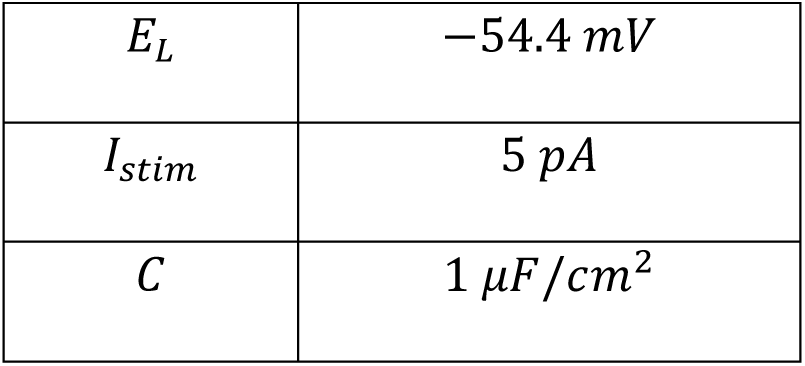
Parameter values used in all simulations

**Table 2.**
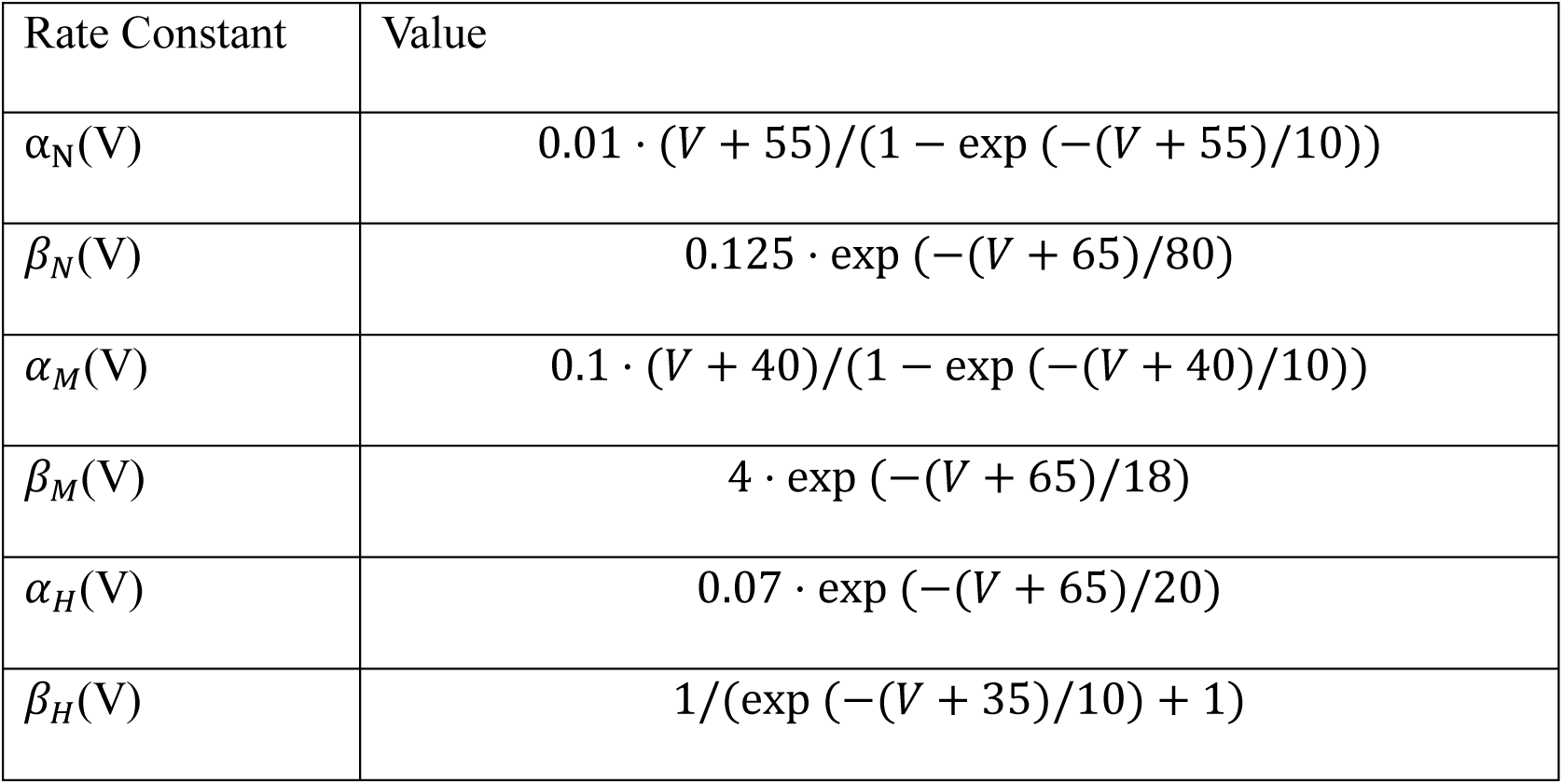
Rate constants of the Hodgkin Huxley model

Generally, the variables (𝑉, 𝑚) have much faster time scales and therefore constitute the fast subsystem of the model, whereas (𝑛, ℎ) constitutes the slower subsystem. This observation becomes particularly important when performing timescale decompositions of the model as will be discussed next.

It is also important to note that the time scale 𝜏_𝑚_(𝑉) is considerably smaller than 𝜏_𝑛_(𝑉) or 𝜏_ℎ_(𝑉). This faster time scale of 𝑚 means that 𝑁𝑎^+^channels activate much more rapidly than they inactivate or than 𝐾^+^ channels open. As a result, there is a brief period of rapid depolarization before the inactivating influence of ℎ or the onset of the potassium repolarizing current. These dynamic interactions create the characteristic spike shape of action potentials. The absolute refractory period that follows each action potential occurs because ℎ remains near zero during the phase of hyperpolarization. During this time, sodium channels remain inactivated, and ℎ must recover and return toward its steady-state value before the neuron can generate another action potential. This recovery process ensures that sodium conductance is temporarily unavailable, preventing immediate re-excitation and ensuring proper signal propagation and temporal separation of action potentials.

### 2.2 Modeling Temperature Effects in Hodgkin-Huxley Dynamics

Ion channels are stochastic in nature and undergo transitions between open and closed states, with energy barriers separating these states. Temperature directly modulates this process by altering the amount of thermal energy available to overcome the barrier, and therefore affects the opening and closing rates of ion channels [3]. Consequently, the gating mechanism of ion channels should be formulated to account for both the voltage and temperature dependence of the gating variables.

As the transition rate constants 𝛼 and 𝛽 of each gating variable best quantify this stochastic channel behavior, Hodgkin and Huxley accounted for the temperature dependence of their computational model by augmenting these rate constants to be functions dependent on both temperature and voltage ( 𝛼(𝑉, 𝑇) and 𝛽(𝑉, 𝑇)). They experimentally determined that the temperature-dependence of these rate constants generally followed the Arrhenius equation [5]. In an Arrhenius-like relation, a temperature-dependent rate constant 𝑥(𝑇) (which, in the context of the Hodgkin-Huxley model, would be a rate constant such as 𝛼_𝑁_(𝑉, 𝑇)) is affected by temperature by the following formula:

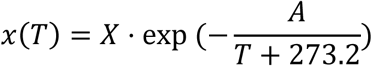

Here, 𝑋 is the pre-exponential factor, the theoretical maximum value the rate constant 𝑥(𝑇) can achieve when there is infinite thermal energy. In the Hodgkin-Huxley model, where rate constants such as 𝛼_𝑁_(𝑉, 𝑇) are highly voltage-dependent, 𝑋 would be a function of voltage. Meanwhile, 𝐴 is the activation energy of the reaction, and 𝑇 is the temperature in degrees Celsius. To simplify this expression, the effect of temperature on the action potential was modelled through a temperature scaling factor 𝑄_10_[19]. It represents the ratio of reaction rates for a 10^∘^𝐶 increase of temperature, or can be thought of as an estimation of rate coefficient increase concerning 10^∘^𝐶 temperature change alteration:

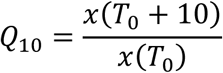

where 𝑇_0_ is the reference or standard temperature.

Substituting this into the Arrhenius equation allows it to be rewritten as:

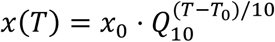

where 𝑥_0_ is the value of 𝑥 at the reference temperature 𝑇_0_. Therefore, temperature can be introduced into the Hodgkin-Huxley model by multiplying each of the rate constants by the temperature dependent factor 𝜙 defined as follows:

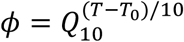

Instead of entirely replacing the notation in Table 2 and redefining each rate constant to be a function dependent on both temperature and voltage, we can simply augment the differential equations of the gating variables to:

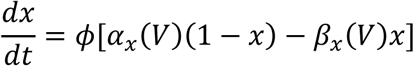

Where 𝑥 = 𝑚, 𝑛 or ℎ, and the rate constants are left in their original, purely voltage-dependent forms (i.e. their values at the default temperature 𝑇_0_). Note this is mathematically equivalent to individually multiplying each rate constant by 𝜙. For this study we will assume this exponential constant to be 𝑄_10_ = 1.5, and set 𝑇_10_ to 25^∘^𝐶 unless otherwise specified.

### 2.3 Bifurcation Analysis

We employed the continuation tool AUTO within XPPAUT to identify regions of excitability and periodic oscillations in our model. The software detects stability transitions and the emergence of stable or unstable limit cycles, thereby facilitating the interpretation of the underlying dynamical behavior. In each two-dimensional projection of the phase space, defined by any pair of state variables (*x*, *y*), the fixed-point continuation yields smooth curves where unstable segments are colored in black and stable segments in red. Transitions in system dynamics frequently occur through Hopf bifurcations (HBs), which are annotated on equilibrium curves. At a Hopf point, a limit cycle is either created or terminated; in the case of a supercritical Hopf bifurcation, the equilibrium loses its stability as a stable limit cycle forms, whereas at a subcritical Hopf bifurcation, an unstable limit cycle collapses into a stable equilibrium, rendering it unstable [42]. Limit cycles are visualized on the bifurcation diagrams as green (stable) and blue (unstable) curves, with their color indicating their stability. In Hodgkin-Huxley type neuronal models, limit cycles often disappear through homoclinic bifurcations (HOMs), where limit cycles collide with saddle points and vanish as their periods diverge to infinity. Another common bifurcation is the limit-point of cycle (LPC) bifurcation, in which a stable and unstable limit cycle coalesce to form a semi-stable cycle at the bifurcation point, which subsequently disappears [42].

Bifurcation analysis of neuronal models reveals that the presence of stable limit cycles corresponds to the ability of neurons to sustain repetitive firing, a hallmark of neuronal excitability. Two primary types of excitability arise from distinct bifurcation mechanisms. Type I excitability typically emerges through LPC bifurcations, where the frequency of the stable limit cycle gradually decreases to zero as the LPC point is approached. Type II excitability emerges via supercritical Hopf bifurcations, in which a stable limit cycle emerges abruptly with a nonzero frequency, resulting in a sudden transition to high-frequency firing [43].

While most of the simulations were carried out using the AUTO and XPPAUT tools, the MATCONT package in MATLAB was also used, primarily to plot two-parameter bifurcation diagrams. Two-parameter bifurcation diagrams are a higher-dimensional generalization of standard bifurcation analysis. In a two-parameter plot, the parameter space is partitioned into regions with different system dynamics (for instance, one region may have bistability with two stable limit cycles), with curves of bifurcation points acting as boundaries between these dynamical regimes. These curves are computed first by locating a codimension-1 bifurcation point with one parameter fixed, then using a two-parameter continuation to generate a smooth curve. Interactions between different curves of bifurcations can create higher-codimension phenomena such as generalized Hopf (GH) bifurcations, at which multiple dynamical regions converge. The inset of Fig. 5 as well as Fig. 6 and Fig. 8 were generated using bifurcation plots in MATCONT. Figure 4 was generated using the standard ODE45 function in MATLAB. All remaining figures were generated by extracting or processing data files from simulations run in XPPAUT, then plotting diagrams as needed in MATLAB using this data.

## Results

We begin by discussing the general codimension-1 bifurcations present in the Hodgkin-Huxley model at various temperatures, and show how these bifurcation diagrams manifest themselves through evolving limit cycles and action potentials. We then provide a comprehensive analysis of how temperature variations influence the model neuron’s dynamics, focusing on the bifurcation phenomena that underlie these transitions.

### General Hodgkin-Huxley Model Dynamics at Standard Temperature (𝑇 = 30^∘^𝐶)

As an initial step, we investigated how solutions of the Hodgkin-Huxley system change when parameters such as the maximal ionic conductances and equilibrium potentials are varied under fixed temperature conditions. Bifurcation theory allows us to classify the qualitative transitions that occur as parameters change, and in particular, to identify the parameter values at which equilibria lose stability and oscillations emerge.

Figure 1 shows the bifurcation diagrams for the main parameters in the model comprising of the maximal conductances (𝑔_𝑁𝑎_, 𝑔_𝐾_, and 𝑔_𝐿_) and the equilibrium potentials (𝐸_𝑁𝑎_, 𝐸_𝐾_ and 𝐸_𝐿_) when temperature is fixed initially at 30°C. The bifurcation structures for the parameters 𝐸_𝐾_ (potassium equilibrium potential) and 𝐸_𝐿_ (leak equilibrium potential) exhibit analogous patterns (Fig. 1A and 1B, respectively). Periodic solution branches for both parameters first appear at a Limit Point of Cycles (LPC) bifurcation, occurring at 𝐸_𝐾_ = −74.63 𝑚𝑉 and 𝐸_𝐿_ = −50.05 𝑚𝑉, where a stable and an unstable limit cycle simultaneously emerge. The unstable limit cycle branch rapidly shrinks in amplitude and collapses into the stable equilibria, as a subcritical Hopf bifurcation occurs at 𝐸_𝐾_ = −71.65 𝑚𝑉 and 𝐸_𝐿_ = −36.26 𝑚𝑉. At these points, two complex conjugate eigenvalues cross the imaginary axis, destabilizing the fixed point and initiating oscillations. For 𝐸_𝐾_, the unstable limit cycle branch undergoes two saddle-node bifurcations before collapsing into the Hopf point, creating a narrow parameter range where multiple unstable limit cycles coexist. Although this has some mathematical significance, these limit cycles do not appear in numerical simulations due to their instability and thus will be generally disregarded in our analysis. The region between the LPC and the first Hopf bifurcation is particularly significant, as it defines a region of bistability where two stable attractors, a stable equilibrium and a stable limit cycle, coexist. Within this range, the system’s behavior depends on the initial stimulus, which can push the neuron into either sustained oscillatory firing or stabilization at resting potential. The limit cycle region (the region at which a stable limit cycle exists) extends far beyond the initial subHopf point. For both parameters, the stable limit cycle branch slowly decreases in amplitude until it eventually collapses into a supercritical Hopf point; this occurs at 𝐸_𝐾_ = −51.13 𝑚𝑉 and 𝐸_𝐿_ = 446 𝑚𝑉, respectively. At the supercritical Hopf point, two conjugate eigenvalues again cross the imaginary line, stabilizing the curve of equilibria. At this point, the equilibrium regains stability as the limit cycle terminates, marking the transition to a monostable regime where the stable equilibrium becomes the sole attractor. In this regime, the neuron always stabilizes at its resting potential, regardless of the initial conditions. For both 𝐸_𝐾_ and 𝐸_𝐿_, the amplitude of the action potentials decreases as a function of the parameter, as illustrated in the distance between the maximum and minimum values on the periodic branches.

**Figure 1.**
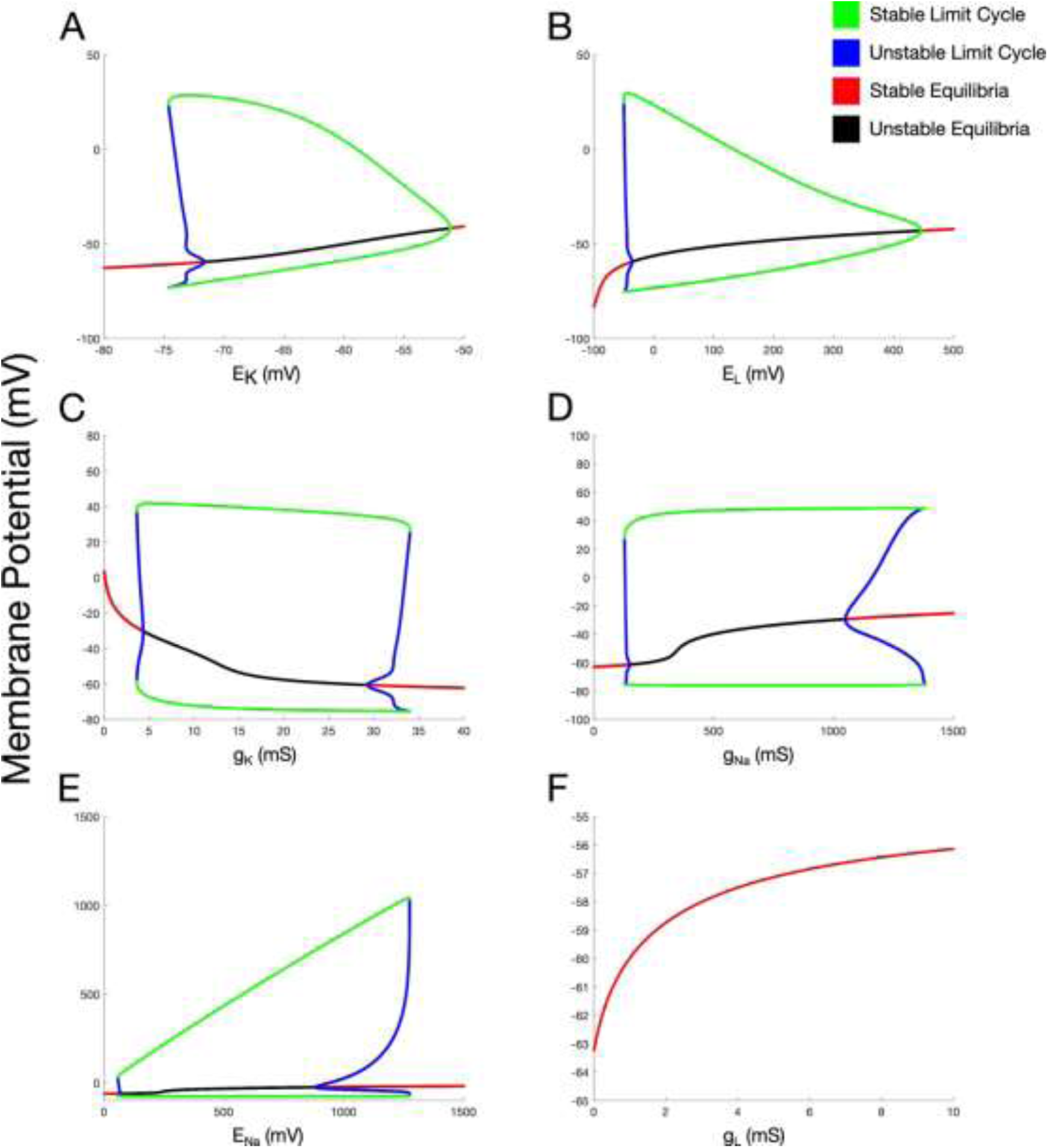
Bifurcation diagrams of the standard parameters, 𝐸_𝐾_(**A**), 𝐸_𝐿_(**B**), 𝑔_𝐾_(**C**), 𝑔_𝑁𝑎_(**D**), 𝐸_𝑁𝑎_(**E**), and 𝑔_𝐿_ (**F**) of the Hodgkin-Huxley model at 30°C. Green and blue lines represent the periodic branches where oscillatory activity is observed. The green lines show the amplitude of the stable limit cycles, while the blue lines indicate unstable limit-cycle phenomena, both born from Hopf bifurcations. The solid red and black lines represent the stationary branches where the stable (red) and unstable (black) fixed points reside.

The potassium and sodium conductances, 𝑔_𝐾_ and 𝑔_𝑁𝑎_, exhibit similar changes in dynamics (Fig. 1C and 1D, respectively). Periodic solutions first emerge at a limit point of cycles (LPC) bifurcation, occurring at 𝑔_𝐾_ = 3.625 𝑚𝑆 and 𝑔_𝑁𝑎_ = 127.2 𝑚𝑆. Subsequntly, the unstable periodic orbit branch merges with the equilibrium curve and destabilizes it at a subcritical Hopf bifurcation at 𝑔_𝐾_ = 4.329 𝑚𝑆 and 𝑔_𝑁𝑎_ = 138.5 𝑚𝑆, defining the first region of bistability between the first LPC and Hopf points. Unlike the other parameters, 𝑔_𝐾_ and 𝑔_𝑁𝑎_ have minimal impact on the amplitude of the stable periodic orbit branch, which is surprising given that these parameters play a significant role in shaping the upstrokes and the downstrokes of the action potentials. A slight decrease in the membrane potential at the peak and trough of stable limit cycles can be observed as 𝑔_𝐾_ increases, whereas 𝑔_𝑁𝑎_ shows no visible effect on the stable orbits. Instead of collapsing at a Hopf bifurcation, the stable periodic solution branch disappears at a second LPC bifurcation where it collides with an unstable limit cycle branch at 𝑔_𝐾_ = 34 𝑚𝑆 and 𝑔_𝑁𝑎_ = 1386 𝑚𝑆. The unstable periodic orbit branch originates from a second subcritical Hopf bifurcation at 𝑔_𝐾_ = 29.29 𝑚𝑆 and 𝑔_𝑁𝑎_ = 1040 𝑚𝑆. At this point, the unstable equilibrium becomes stabilized as an unstable limit cycle is born around it. This establishes a second region of bistability between the second Hopf and LPC bifurcations, where a stable equilibrium and stable limit cycle coexist. Thus, for the parameters 𝑔_𝐾_ and 𝑔_𝑁𝑎_, there are two disjoint regions of bistability present. Beyond the final LPC bifurcation, the system transitions into a region of monostable dynamics, with the stable equilibrium acting as the unique attractor.

The sodium Nernst potential (𝐸_𝑁𝑎_) exhibits a bifurcation structure intermediate between the two types discussed earlier (Fig. 1E). As with the other parameters, periodic solutions emerge from a limit point of cycles (LPC) bifurcation at 𝐸_𝑁𝑎_ = 56.79 𝑚𝑉. Subsequently, the unstable limit cycle branch merges with the stable fixed point, destabilizing it at a subcritical Hopf bifurcation occurring at 𝐸_𝑁𝑎_ = 76.33 𝑚𝑉. The interval between the first LPC and the first Hopf point, 56.79 𝑚𝑉 < 𝐸_𝑁𝑎_ < 76.33 𝑚𝑉, defines the first region of bistability. In contrast to 𝑔_𝐾_ and 𝑔_𝑁𝑎_, 𝐸_𝑁𝑎_ has a pronounced effect on the amplitude of the limit cycles (and thereby the amplitude of the action potentials). As 𝐸_𝑁𝑎_ increases within the excitability region, the peak membrane potential of the limit cycles rises sharply, approaching +1200 𝑚𝑉 while the trough remains relatively constant at around −40 𝑚𝑉. At 𝐸_𝑁𝑎_ = 1273 𝑚𝑉, however, the stable limit cycle branch coalesces with an unstable cycle branch and disappears at a second LPC point. The unstable limit cycle branch bifurcates out of another subcritical Hopf bifurcation at 𝐸_𝑁𝑎_ = 881.1 𝑚𝑉, defining a second region of bistability in the interval, 881.1 𝑚𝑉 < 𝐸_𝑁𝑎_ < 1273 𝑚𝑉. Beyond the second LPC point, the system is monostable, with a single stable fixed point. Finally, the maximal leak conductance (𝑔_𝐿_) does not exhibit any evident dynamical shifts in the region of the parameter space in question. Increasing 𝑔_𝐿_, however, simply causes a rise in the membrane potential at which the neuron stabilizes (the stable equilibrium), reflecting how the leak current has the effect of drawing the membrane potential near 𝐸_𝐿_ (−54.4 𝑚𝑉).

In addition to the maximal conductances and the equilibrium potentials, the effects of the external applied current on the morphological properties of the action potentials and the general dynamics of the HH model is illustrated in Figure 2 in more granular details. Fig. 2A shows a numerical bifurcation diagram of the model in response to constant current injection (𝐼_𝑠𝑡𝑖𝑚_), illustrating how oscillations can emerge in a Hopf bifurcation with increasing drive. In particular, periodic solutions first appear at an LPC bifurcation point at 𝐼_𝑠𝑡𝑖𝑚_ = 6.304 𝑝𝐴. The unstable orbit branch collapses into a subcritical Hopf bifurcation point occurring at 𝐼_𝑠𝑡𝑖𝑚_ = 10.44 𝑝𝐴, while the stable orbit branch more gradually shrinks in amplitude and terminates at a supercritical Hopf point at 𝐼_𝑠𝑡𝑖𝑚_ = 155.1 𝑝𝐴. The interval between the LPC and subcritical Hopf bifurcation (6.304 𝑝𝐴 < 𝐼_𝑠𝑡𝑖𝑚_ < 10.44 𝑝𝐴) determines the region of bistability for the neuron.

**Figure 2.**
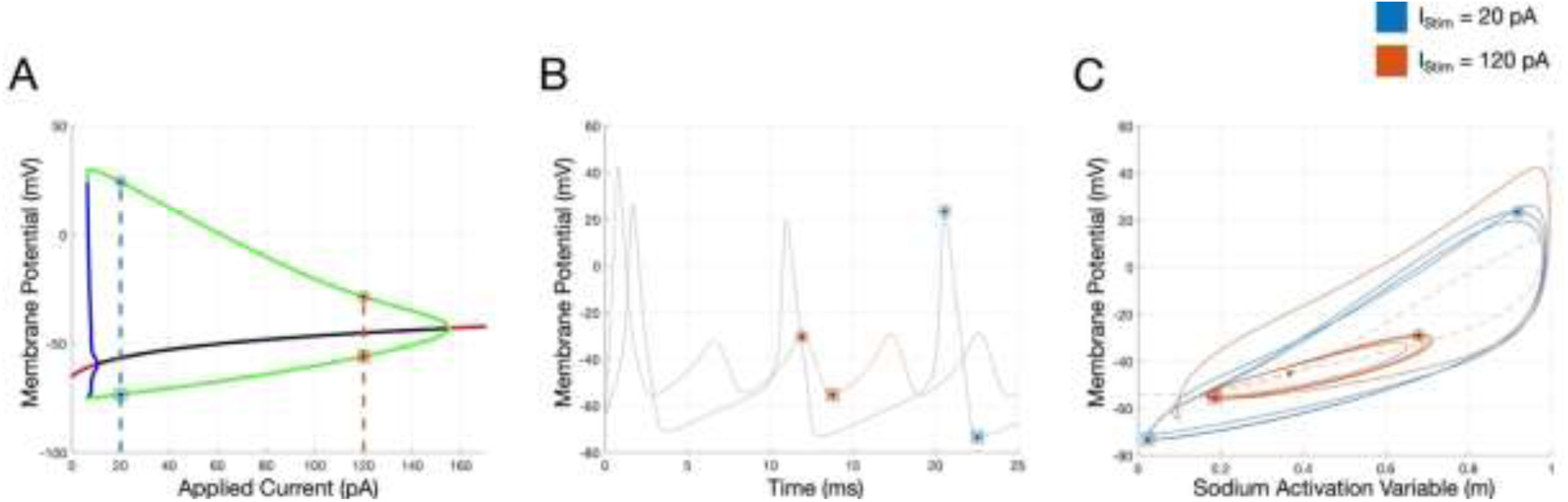
Effects of the external drive on the neuron’s dynamics. **A.** Bifurcation diagram of the HH model as a function of the applied current (𝐼_𝑠𝑡𝑖𝑚_). Panel is color coded as in Fig. 1. Blue (for 𝐼_𝑠𝑡𝑖𝑚_ = 20 pA) and red (for 𝐼_𝑠𝑡𝑖𝑚_ = 120 pA) dashed lines intersect with the periodic branches at the maximum/peak voltage (circles), as well as the minimum/nadir voltage (squares) elicited for the corresponding action potentials. **B.** Membrane potentials as a function of time for the two magnitudes of applied current selected in **A**. The peak (circles) and nadir (squares) voltage values for the third action potentials elicited in response to 20 pA (blue) and 120 pA (red) applied currents are shown, illustrating the difference in spike amplitude and other morphologies. **C.** Overlaid trajectories in the 𝑉 − 𝑚 plane at the two 𝐼_𝑠𝑡𝑖𝑚_ values (blue for 20 𝑝𝐴 and red for 120 𝑝𝐴) beginning from the same initial conditions (𝑉, 𝑚, 𝑛, ℎ) = (−65,0.1,0.1,0.1). The *V-* and *m-* nullclines intersect between the two knees of the cubic *V-* nullcline at unstable steady states ( (𝑉, 𝑚, 𝑛, ℎ) = (−56.594,0.13457,0.45046,0.30768) when 𝐼_𝑠𝑡𝑖𝑚_ = 20𝑝𝐴 and (𝑉, 𝑚, 𝑛, ℎ) = (−45.122,0.36609,0.61748,0.08860) when 𝐼_𝑠𝑡𝑖𝑚_ = 120𝑝𝐴) that coexist with a stable limit cycle. The nullclines of the system when 𝐼_𝑠𝑡𝑖𝑚_ = 120𝑝𝐴 are shown as red dotted curves, intersecting at the unstable equilibrium marked with a red dot. The limit cycle spans a much smaller interval of membrane potential (for 120 pA) compared to that for 20 pA.

Increasing the magnitude of the positive applied current has a negative effect on the amplitude or peak of the action potentials in the HH model (and this mechanism is similar to that of 𝐸_𝐾_ and 𝐸_𝐿_). This can be illustrated by selecting two different magnitudes of 𝐼_𝑠𝑡𝑖𝑚_(20 pA and 120 pA) and checking the corresponding membrane trajectories in the phase plane plots (Fig. 2B-2C). For example, while both magnitudes of applied current generate sustained oscillations, for an external drive of 20 pA (the blue dashed line in Fig 2A), the action potential amplitude shown in Fig. 2B (blue trace) is larger than that elicited for an external drive of 120 pA (red dashed line in Fig. 2A, and red trace in Fig. 2B). In all panels (Fig. 2A-2C), circles denote the maximum voltage elicited for the corresponding magnitude of applied current (blue for 20 pA and red for 120 pA), while squares denote the minimum voltage elicited.

This can be explained further by overlaying the trajectories (limit cycles) of both spike trains in the *V-m* phase plane plot (Fig. 2C). The *V-* and *m-* nullclines intersect between the two knees of the cubic *V-* nullcline at unstable steady states ( (𝑉, 𝑚, 𝑛, ℎ) = (−56.59,0.13457,0.45046,0.30768) when 𝐼_𝑠𝑡𝑖𝑚_ = 20𝑝𝐴 and (𝑉, 𝑚, 𝑛, ℎ) = (−45.122,0.36609,0.61748,0.08860) when 𝐼_𝑠𝑡𝑖𝑚_ = 120𝑝𝐴) (Fig 2C), which coexist with a stable limit cycle. In short, when a positive external current is applied, the low-voltage portion of the *V-* nullcline moves up whilst the high-voltage part remains relatively unchanged. For sufficiently large constant external input, the intersection of the two nullclines falls within the two knees of the cubic *V-* nullcline. In this case, the fixed point is unstable and the emergent limit cycle is stable, producing a train of action potentials and rendering the system as being oscillatory. For 𝐼_𝑠𝑡𝑖𝑚_ of 120 pA, the limit cycle spans a much smaller interval of membrane potential compared to the blue cycle (for 𝐼_𝑠𝑡𝑖𝑚_ of 20 pA). This is primarily due to the increased excitability that the model neuron undergoes when stimulated by larger applied currents. Lower-amplitude stable limit cycles correspond to lower-amplitude action potential chains, and this comes at a tradeoff with higher frequency of action potentials: although this relationship is far from rigorous, we speculate that lower-amplitude action potential chains occur at higher firing frequencies as there is less variation in the gating variables (particularly the sodium inactivation variable ℎ) and thus shorter refractory periods. Therefore, although lower-amplitude stable limit cycles correspond to weaker action potentials, they suggest an increased firing rate. Potential implications of these changes include reduced synaptic transmission efficiency due to weaker simulation of the postsynaptic cell, or higher metabolic efficiency (due to weaker action potentials) with the tradeoff of compromised signal reliability.

### Influence of Temperature on Model Dynamics

The effects of temperature on the HH model neuron are next investigated by studying the changes in the various intrinsic properties in response to temperature perturbations (Fig. 3 and 4). The spiking time patterns at three different temperatures (20^∘^𝐶, 45^∘^𝐶 and 70^∘^𝐶) are illustrated in Fig. 3A where the external applied current was kept constant at 10 pA. At 20^∘^𝐶 (pink trace), the model neuron exhibits lower firing frequency, and the corresponding action potentials exhibit larger amplitudes, wider widths and longer delays to spiking than those elicited at 45^∘^𝐶 (reddish brown trace). At larger magnitudes of temperature (for example, 70^∘^𝐶, purple trace), the model neuron halts from generating sustained firing. This can be better illustrated in the inset of Fig. 3A that depicts the firing frequency of the model neuron as a function of increasing temperature. The number of spikes increases almost linearly as temperature is raised, but when temperature reaches a value of 66.39^∘^𝐶 (for 10 pA applied current), the model HH neuron halts from firing entirely.

**Figure 3.**
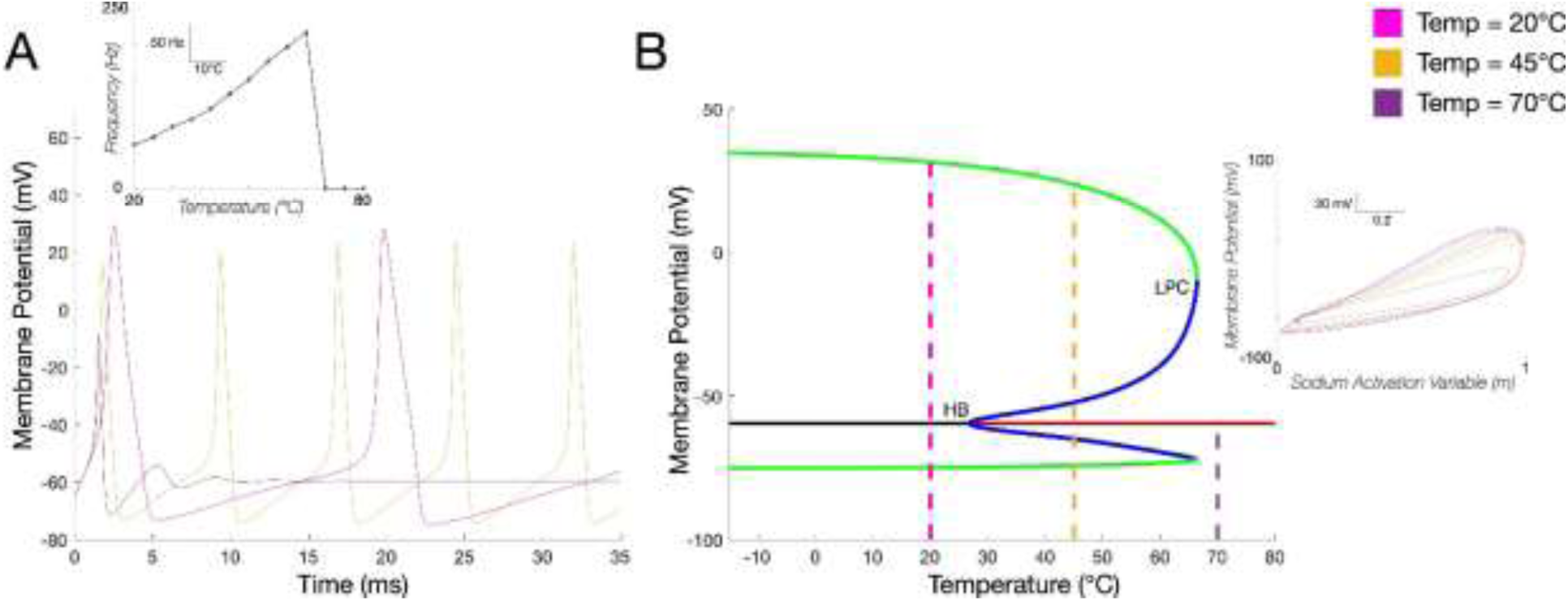
Effects of temperature on the firing frequency and bifurcation diagram of HH model neuron. **A.** Firing patterns of the model HH neuron at temperatures of 20^∘^𝐶 (pink), 45^∘^𝐶 (reddish brown), and 70^∘^𝐶 (purple), under the same initial conditions (𝑉, 𝑚, 𝑛, ℎ) = (−65,0.1,0.1,0.1) and for the same magnitude of applied current (10 pA). As temperature increases, spike widths narrow, spike amplitudes shorten, and the delay to spiking decrease. *Inset*: Firing frequency increases almost linearly as a function of temperature. At T = 66.39^∘^𝐶, model neuron stops firing entirely. **B**. Bifurcation diagram with temperature as the bifurcation parameter. Subcritical HB occurs at 26.79^∘^𝐶, and before it a stable limit cycle persists throughout low temperature ranges, terminating at an LPC bifurcation point at 66.39^∘^𝐶. A bistable region occurs for temperature values between 26.79^∘^𝐶 and 66.39^∘^𝐶. Dashed lines illustrate the values in temperature on the bifurcation diagram where the various simulations are shown in panel **A**. *Inset*: Phase plane trajectories in the (𝑉 − 𝑚) subsystem under the three different temperatures.

Fig. 3B shows the bifurcation diagram as temperature is varied. A stable periodic orbit branch persists throughout low temperature ranges, and terminates at an LPC bifurcation point at 66.39^∘^𝐶. The unstable limit cycle branch with which the stable orbit branch collides originates from a subcritical Hopf bifurcation at 26.79^∘^𝐶. At 20^∘^𝐶 the model is in the region before the Hopf bifurcation (𝑇𝑒𝑚𝑝 < 26.79^∘^𝐶), and thus the only stable attractor is a limit cycle. This can be seen in the inset of Fig. 3B, where the trajectory in pink converges to a periodic orbit, representing sustained firing where the model neuron is highly excitable, always reaching a sustainable and continuous chain of nerve impulses in response to a presynaptic stimulus regardless of its initial conditions. At 45^∘^𝐶, the model is in the region of bistability, 26.79^∘^𝐶 < 𝑇𝑒𝑚𝑝 < 66.39^∘^𝐶, as a stable limit cycle and stable steady state coexist in this regime. Accordingly, some trajectories converge to the stable limit cycle, whereas some other trajectories converge to the stable equilibrium. This means that the neuron is at a level of excitability where minuscule changes in its stimulus could cause it to either exhibit perpetual nerve impulse chains or dampen its excitability and stabilize it at resting membrane potentials. As noted previously, in this region the neuron is particularly sensitive to changes in the presynaptic stimulus. When 𝑇𝑒𝑚𝑝 = 70^∘^𝐶, the model is in the region of monostability (𝑇𝑒𝑚𝑝 > 66.39^∘^𝐶) where a stable equilibrium acts as the unique attractor after the stable limit cycle is destroyed through the LPC bifurcation. Note that, when approaching a stable equilibrium, the trajectory spirals inward. This reflects the successive action potentials gradually dampening until the neuron stabilizes at resting potential.

Moreover, the trajectories in the (𝑉, 𝑚) phase plane (the inset of Figure 3B) show that as temperature increases, the distance spanned in the voltage domain gets narrower (indicating shorter spikes), the peak of Na^+^ activation (𝑚) gets smaller (indicating faster activation kinetics), and the time it reaches peak Na^+^ activation is faster (indicated by shallower slopes on the 𝑚 axis). As noted in Fitzhugh (1966), there are two central mechanisms through which temperature affects the action potential morphology of neurons. First, the timescales of the three gating variables 𝑚, 𝑛, and ℎ decrease exponentially as temperature increases, due to the 𝑄_10_ factor in the differential equations. Faster 𝑚 kinetics means that the initial excitation process occurs much more rapidly, while faster 𝑛 and ℎ kinetics allow for a more rapid depolarization and shorter absolute refractory period.

Overall, the faster evolution of the gating variables expedites the action potential process, decreasing its period. In general, the period of the stable limit cycle branch decreases seemingly exponentially as temperature rises. This has the effect of making the action potentials ’thinner’ in the 𝑉 − 𝑡 plane, indicating overall a shorter action potential allowing for a higher firing rate. The second effect is that, at sufficiently high temperatures, the timescale decomposition between the subsystems (𝑉, 𝑚) and (𝑛, ℎ) break down as the timescales of the gating variables approach that of 𝑉. This is because, whereas the three gating variables speed up exponentially as temperature rises, the dynamics of 𝑉 remains relatively unaffected. An interesting observation is that the position of the equilibrium point is seemingly unaffected by temperature. For instance, at the subcritical Hopf point an unstable limit cycle is formed around the equilibrium as it gains stability, but the position of the equilibrium point never changes. The reason for that is because equilibrium points in the HH model are points (𝑉^∗^, 𝑚^∗^, 𝑛^∗^, ℎ^∗^) satisfying the set of equations (𝑉*^′^*, 𝑚*^′^*, 𝑛*^′^*, ℎ*^′^*) = (0,0,0,0). As temperature simply scales the values of 𝑚*′*, 𝑛*′*, and ℎ*′* by a nonzero exponential factor, the set of solutions to these equations are unaffected by temperature. Thus, the equilibria remain invariant as well.

Temperature fluctuations have detrimental effects on the intrinsic properties of model HH neurons (Fig. 4). As shown, an exponential decrease in spike width occurs as temperature is increased (Fig. 4A), and its inset shows the first spike morphologies from the three spike trains of Fig. 3A, illustrating the apparent increase in spike width under the three different temperatures. Similar effects of increasing temperature values are observed on spike threshold (Fig. 4C), delay to first spike from current onset (Fig. 4D), time to action potential peak from the threshold point (Fig. 4E), and the after-hyperpolarization magnitude (Fig. 4F). Spike amplitude has a slightly different dynamics, where it increases a little as temperature is increased and momentarily plateaus near 40^∘^𝐶 before it undergoes a rapid decline (Fig. 4B). In short, as temperature increases, the inactivating gating variable ℎ reaches its steady state and blocks the depolarizing sodium current before membrane potential reaches its usual peak value. This is responsible for the sharp decline in action potential amplitude as temperature increases beyond a certain range (Fig. 3B and 4B), as well as the decrease in spike width (Fig. 4A), especially in the region after the HB. The decrease in amplitude is also partly due to the fact that the repolarizing potassium current is established quickly due to faster 𝑛 dynamics, providing a smaller window for the neuron to freely depolarize. The second effect is primarily responsible for the lack of a stable periodic orbit beyond the final LPC bifurcation point. At sufficiently high temperatures, 𝑉 can be considered as the slow subsystem and (𝑚, 𝑛, ℎ) the fast subsystem of the Hodgkin-Huxley model, and thus at the slow time scale the model becomes of the form 𝑉*^′^* = 𝑓(𝑉) where 𝑓 is a function solely dependent on 𝑉. As this is a one-dimensional flow, it is impossible for the system to generate periodic solutions, rendering the neuron incapable of firing continuous action potentials.

**Figure 4.**
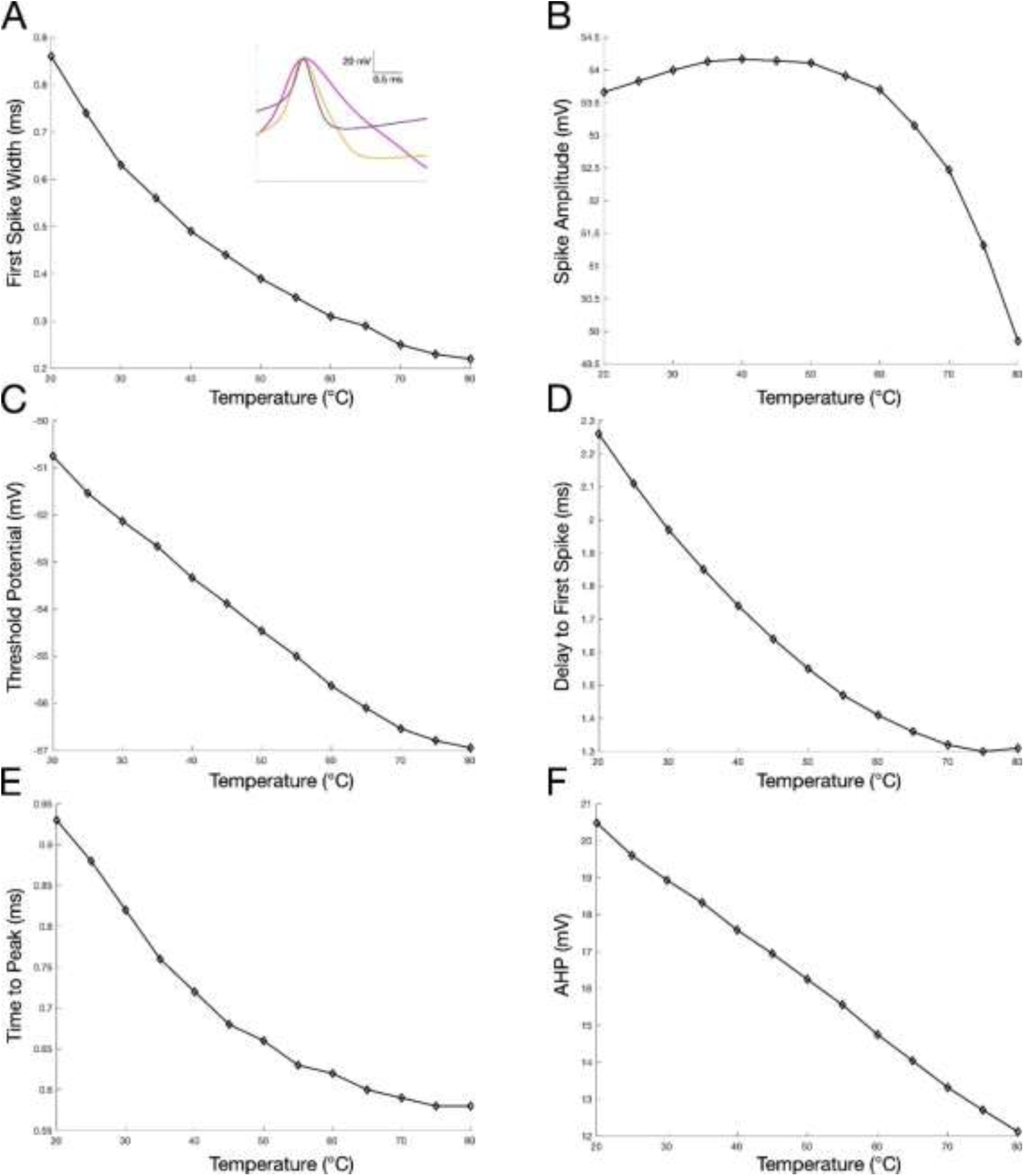
Effects of temperature on the intrinsic properties of HH model neuron. **A.** The spike widths of action potentials decrease exponentially as temperature is increased. *Inset*: Action potentials morphologies for the first spikes in the spike trains in (Fig. 3A) depicting the decrease in spike width as temperature is increased. Similar effects of increasing temperature values are observed on spike amplitude (**B**), spike threshold (**C**), delay to first spike from current onset (**D**), time to action potential peak from the threshold point (**E**), and the after-hyperpolarization magnitude (**F**).

Therefore, increasing temperature increases the firing rates at the cost of weaker action potentials (Fig. 3 and Fig. 4), which may expedite signal transmission but also may, in extreme cases, inhibit the presynaptic neuron from providing a strong enough stimulus to activate successive postsynaptic action potentials.

### Generalized Hopf (Bautin) Bifurcations Involving Temperature

Changes in the applied current switches the dynamics from stationary to periodic on the bifurcation diagram as shown earlier (Fig. 2A); however, temperature fluctuations can distort the shape and various topological properties of the bifurcation diagram, which is eventually a direct reflection of the changes in the firing patterns of the underlying neurons. To better visualize this, we overlaid the bifurcation diagrams of 𝐼_𝑠𝑡𝑖𝑚_ at four different temperature values (Fig. 5): 20^∘^𝐶 (purple), 40^∘^𝐶 (yellow), 60^∘^𝐶 (blue), and 80^∘^𝐶 (green). As temperature rises, stronger external applied currents are needed to start triggering action potentials, the interval of oscillations (sustained firing) diminishes, and the amplitudes of the action potentials shorten. While each bifurcation diagram exhibits a subcritical and supercritical Hopf bifurcation as temperature is varied, the distance between the two (in the 𝐼_𝑠𝑡𝑖𝑚_ space) is diminished as temperature is increased, highlighting smaller regions of excitability. The rightward movement of the first LPC point and the leftward shift of the second Hopf point causes the limit cycle region to be squeezed from both ends. In fact, as temperature rises from 20^∘^𝐶 to 80^∘^𝐶, the limit cycle region contracts substantially from 6.245 𝑝𝐴 < 𝐼_𝑠𝑡𝑖𝑚_ < 153.9 𝑝𝐴 to 20.01 𝑝𝐴 < 𝐼_𝑠𝑡𝑖𝑚_ < 117.2 𝑝𝐴. This is illustrated in Fig. 5 where the distance between every square (representing subcritical HB) and circle (representing supercritical HB) of the same color (representing a particular temperature) gets smaller and smaller as a function of temperature. This is further illustrated in the insets (A and B) of Fig. 5 where the values of the subcritical (red) and supercritical (blue) Hopf bifurcations are plotted for a wider range of temperatures. Notice that the differences between the two Hopf points whether in their voltage domain (A) or their current domain (B) get smaller and smaller as temperature increases, eventually coalescing.

**Figure 5.**
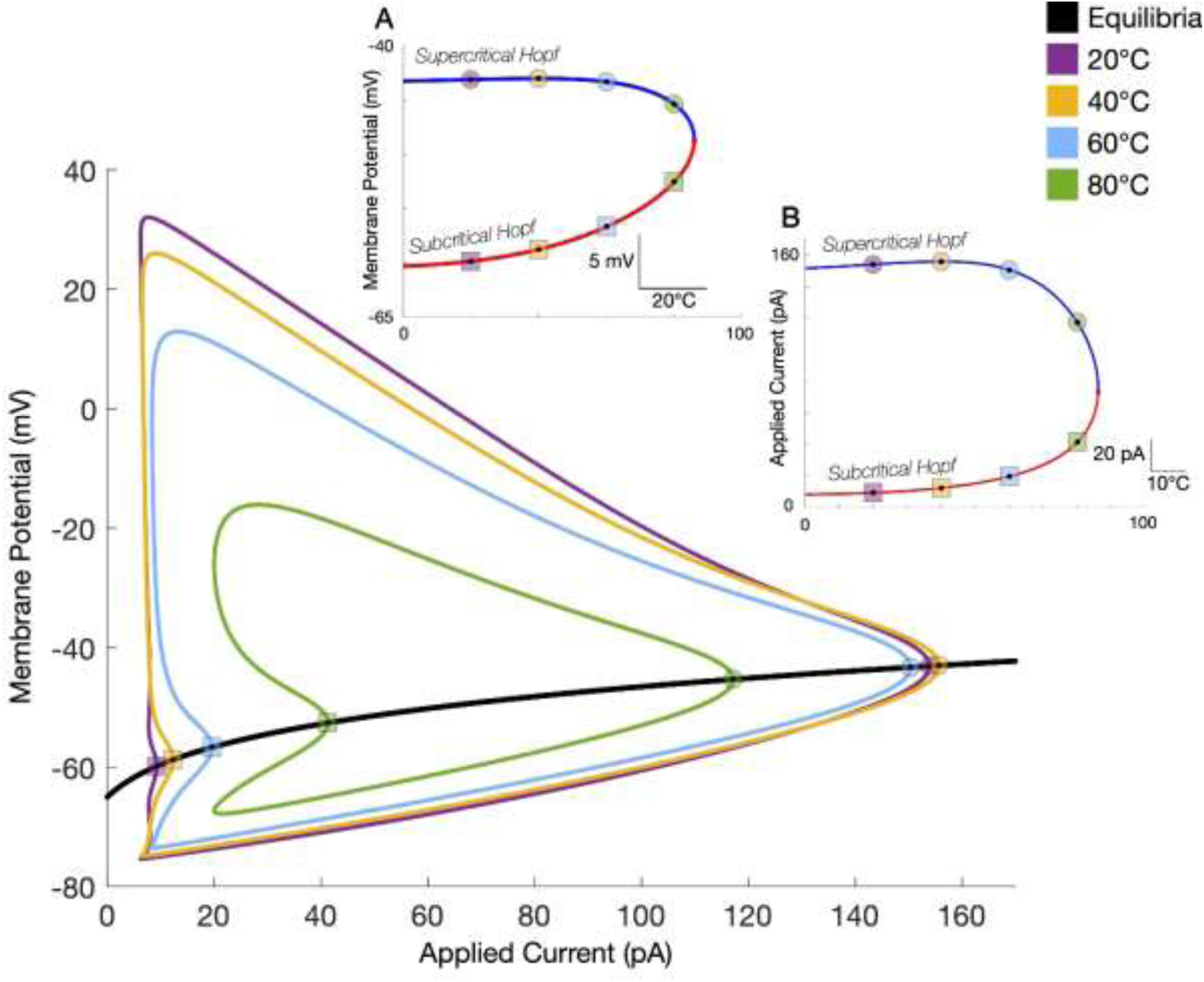
An overlay of bifurcation diagrams of applied current at different temperature values (20^∘^𝐶 (purple), 40^∘^𝐶 (yellow), 60^∘^𝐶 (blue), and 80^∘^𝐶 (green)). The curve of equilibria (black) does not change as temperature varies; it is only the position of bifurcations and limit cycles that are affected. Squares represent subcritical Hopf points and circles represent supercritical Hopf points. The distance between the two gets smaller and smaller as temperature increases. *Inset* **(A-B)**: Values of the subcritical (red curve) and supercritical (blue curve) Hopf bifurcations are plotted for a wider range of temperatures. The differences between the two Hopf points whether in their voltage domain (**A**) or their current domain (**B**) gets smaller and smaller as temperature increases, eventually coalescing at a temperature of 86.1062^∘^𝐶 (𝐼_𝑠𝑡𝑖𝑚_ = 73.0902 pA and 𝑉 = -48.8314 mV at the coalesced Hopf point).

**Figure 6.**
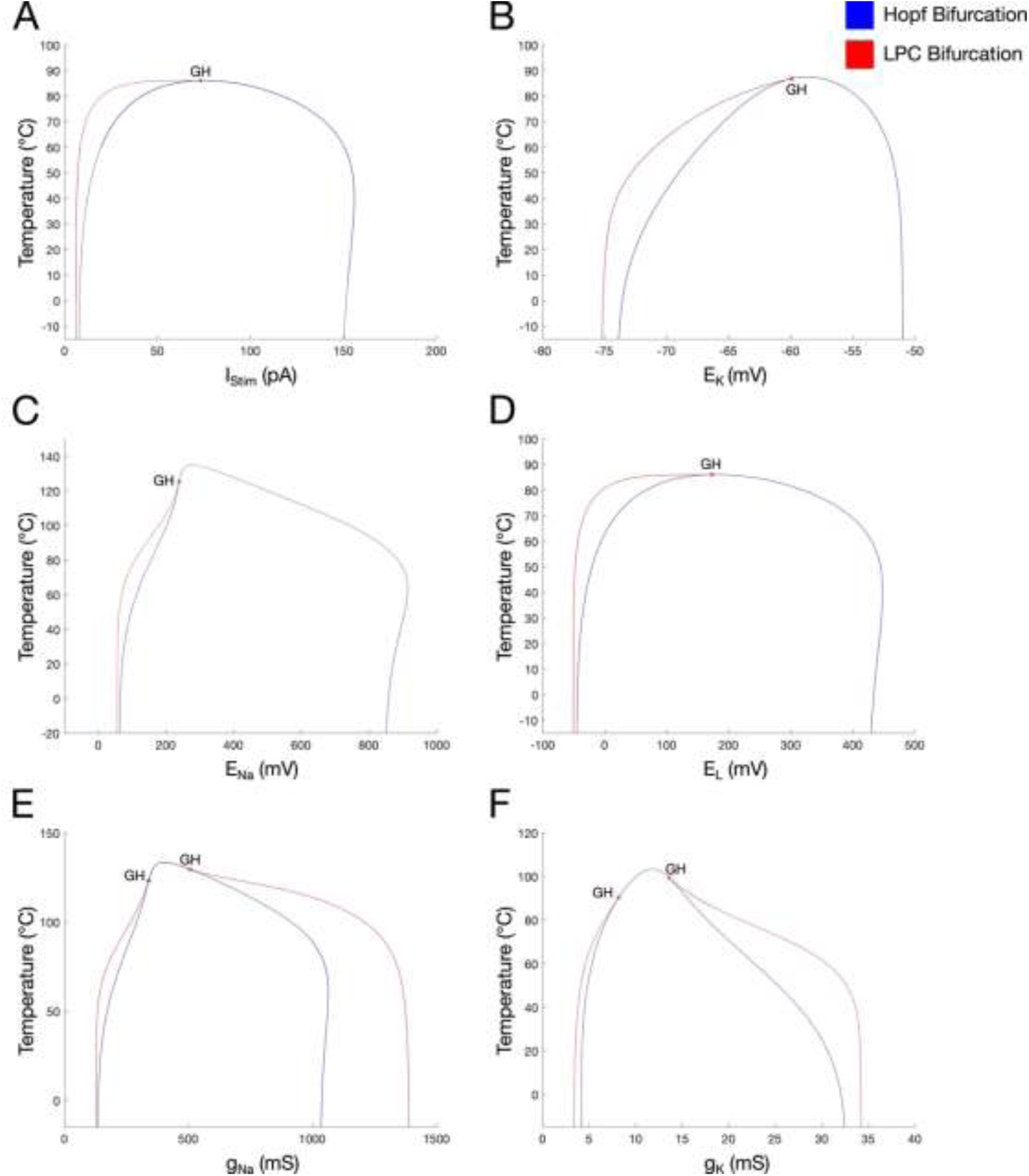
Two-parameter bifurcation diagrams for standard system parameters, 𝐼_𝑠𝑡𝑖𝑚_ (**A**), 𝐸_𝐾_ (**B**), 𝐸_𝑁𝑎_(**C**), 𝐸_𝐿_(**D**), 𝑔_𝑁𝑎_(**E**), and 𝑔_𝐾_(**F**) with temperature. The blue curve represents Hopf bifurcation points whereas the red denotes a curve of LPC points. GH denotes generalized Hopf (or Bautin) bifurcation points.

To explain the mechanisms behind the bifurcation diagrams briefly, as temperature increases, the ℎ gating kinetics becomes faster, facilitating the inhibition of action potential generation; in other words, the weakened depolarization mechanism due to faster sodium inhibition necessitates stronger external applied currents to trigger trains of action potentials. Additionally, the earlier onset of repolarization due to accelerated 𝑛 gating kinetics weakens the excitability of the neuron. This is evidenced in Fig. 5 by the 𝐼_𝑠𝑡𝑖𝑚_threshold for sustained firing (the first LPC point) increasing from 6.245 𝑝𝐴 to 20.01 𝑝𝐴 as temperature increases from 20^∘^𝐶 to 80^∘^𝐶. Moreover, the inhibited dynamics of membrane potential due to the dissolving timescale decomposition at high temperatures exacerbates the depolarization block phenomenon, where the neuron is unable to fire at high 𝐼_𝑠𝑡𝑖𝑚_values. The depolarization block occurs due to the neuron remaining perpetually depolarized; this prevents ℎ from recovering and allowing for a further depolarizing sodium current, inhibiting the onset of an action potential. At higher temperatures, the relatively slow evolution of 𝑉 makes the depolarization block occur at lower 𝐼_𝑠𝑡𝑖𝑚_ values due to the fact that sufficiently high temperatures assist the depolarization block phenomenon in preventing perpetual firing chains. This is evidenced in Fig. 5 where the second Hopf point (where depolarization block begins to occur) occurs at a lower 𝐼_𝑠𝑡𝑖𝑚_ value of 117.2 𝑝𝐴 at 80^∘^𝐶 compared to 153.9 𝑝𝐴 at 20^∘^𝐶. Moreover, the curvature of the unstable limit cycle curve intermediating the LPC and first Hopf points increases as temperature increases. This has the effect of expanding the region of bistability; the bistability region increases from 6.245 𝑝𝐴 < 𝐼_𝑠𝑡𝑖𝑚_ < 9.254 𝑝𝐴 at 20^∘^𝐶 to 20.01 𝑝𝐴 < 𝐼_𝑠𝑡𝑖𝑚_ < 41.3 𝑝𝐴 at 80^∘^𝐶. Physiologically speaking, this indicates neurons are more signal-sensitive in terms of nerve impulse firing patterns at high temperatures, as they are more likely to be at the region of bistability.

As mentioned earlier, the limit cycle region contracts substantially as temperature increases. It is reasonable to expect this contraction to continue until periodic orbits are no longer obtainable regardless of the external current applied. This was shown in Fig. 3, in which the stable limit cycle disappears at temperatures beyond the LPC bifurcation point. In fact, this phenomenon is better explained through two-parameter bifurcation diagrams as illustrated in Fig. 6. The limit cycle region for 𝐼_𝑠𝑡𝑖𝑚_shrinks further and further until it disappears at a generalized Hopf (or Bautin) bifurcation point at 86.1^∘^𝐶 and 𝐼_𝑠𝑡𝑖𝑚_of 73.1 𝑝𝐴 (Fig 6A); this can be analytically verified by the fact that the first Lyapunov coefficient vanishes at this point on the Hopf curve. At temperatures higher than 86.1^∘^𝐶, the system consistently displays monostability with a unique stable equilibrium point as there are no further Hopf bifurcations to generate limit cycles or switch the stability of equilibria. The type of behavior and the general shifts in dynamics due to temperature fluctuations are analogous across all parameters, where the limit cycle region gradually shrinks until disappearing entirely at a generalized Hopf bifurcation point as temperature increases. Beyond a particular critical temperature threshold, the system becomes monostable with a unique stable equilibrium point. A notable distinction to make however is between parameters with one and those with two Bautin bifurcation points. For example, besides the 𝐼_𝑠𝑡𝑖𝑚_parameter described above, 𝐸_𝐾_(Fig. 6B), 𝐸_𝑁𝑎_(Fig. 6C) and 𝐸_𝐿_(Fig. 6D) all exhibit one LPC bifurcation branch and one region of bistability, where the LPC branch coalesces with the supercritical and subcritical Hopf branches at a single generalized Hopf point. On the other hand, the 𝑔_𝑁𝑎_ (Fig. 6E) and 𝑔_𝐾_ (Fig. 6F) parameters exhibit two subcritical Hopf bifurcations leading into two different LPC bifurcations that have two distinct Bautin points at which the branches terminate, with each LPC point curve ending at separate bifurcations.

### Generalizing Dynamical Effects of Timescale Warping

There are several different methods of incorporating temperature into the Hodgkin-Huxley model, each with different dynamical behavior and bifurcations. Although we used 𝑄_10_ = 1.5 in our previous analysis, Hodgkin and Huxley assumed 𝑄_10_ to be 3 in their original paper, while Fitzhugh (1986) also multiplied the channel conductances (𝑔_𝐾_, 𝑔_𝑁𝑎_, and 𝑔_𝐿_) by a linear factor 𝜈 = 𝐴[1 + 𝐵(𝑇 − 6.3)]. Most models involve some manner of multiplying the gating variable transition constants by an exponential factor. The exponential factor by which each gating variable is warped does not necessarily have to be equal as well; for instance, 𝑛 may have a 𝑄_10_ factor of 1.5 whereas ℎ has a factor of 2. We will expand our analysis to include all such types of models.

To begin, we formulated a model with three additional parameters to generalize this temperature-dependence. This can be done by multiplying the transition rates of the three gating variables with scalar factors 𝑥, 𝑦, and 𝑧, as follows:

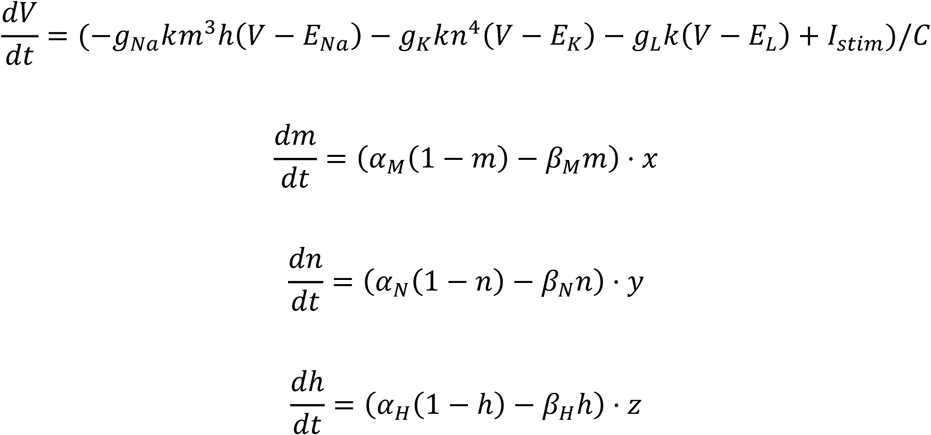

Note that the model used for our previous analysis is a special case where 𝑥 = 𝑦 = 𝑧 = (1.5)^(𝑇𝑒𝑚𝑝−25)/10^. Furthermore, as we wish our analysis to be realistically reproducible, we will limit the parameters (𝑥, 𝑦, 𝑧) to the interval (0.1,10).

Figure 7 provides an overview of how shifts in the scaling factors 𝑥, 𝑦, and 𝑧 change the dynamics of the model and alter the action potentials of the neurons as depicted on the underlying bifurcation diagrams. First, for the 𝑥 scaling factor (Fig. 7A1), the system displays monostable dynamics at low values with a single stable equilibrium point until a stable limit cycle appears at an LPC bifurcation, occurring at 𝑥 = 0.5432. The unstable limit cycle branch collapses into the stable equilibrium, destabilizing the equilibrium at a subcritical Hopf bifurcation at 𝑥 = 0.9355. At 𝑥 > 0.9355, the system is again monostable with a limit cycle being the unique attractor. Note that the interval 0.5432 < 𝑥 < 0.9355 is the region of bistability, where the neuron is particularly sensitive to external stimulus as its ultimate behavior is dependent on the initial conditions. Fig. 7A2 provides more insights into why these bifurcations occur, where the membrane potential trajectories over time are overlaid for three different values of 𝑥 (0.5 in pink, 1 in yellow and 2 in purple – same values are selected in Fig. 7B2 and 7C2). Increasing 𝑥 expedites the depolarization mechanism through fast sodium activation, creating higher-amplitude spikes with more precipitous depolarization curves. In contrast, lower 𝑥 values can impede action potentials (as demonstrated by the 𝑥 = 0.5 curve), reflected by the spontaneous disappearance of the stable limit cycle branch at the LPC bifurcation at 𝑥 = 0.5432. Biologically, increasing the m-scaling variable makes the neuron more excitable and its action potentials larger, facilitating synaptic transmission by allowing the presynaptic neuron to more easily stimulate its postsynaptic counterpart.

**Figure 7.**
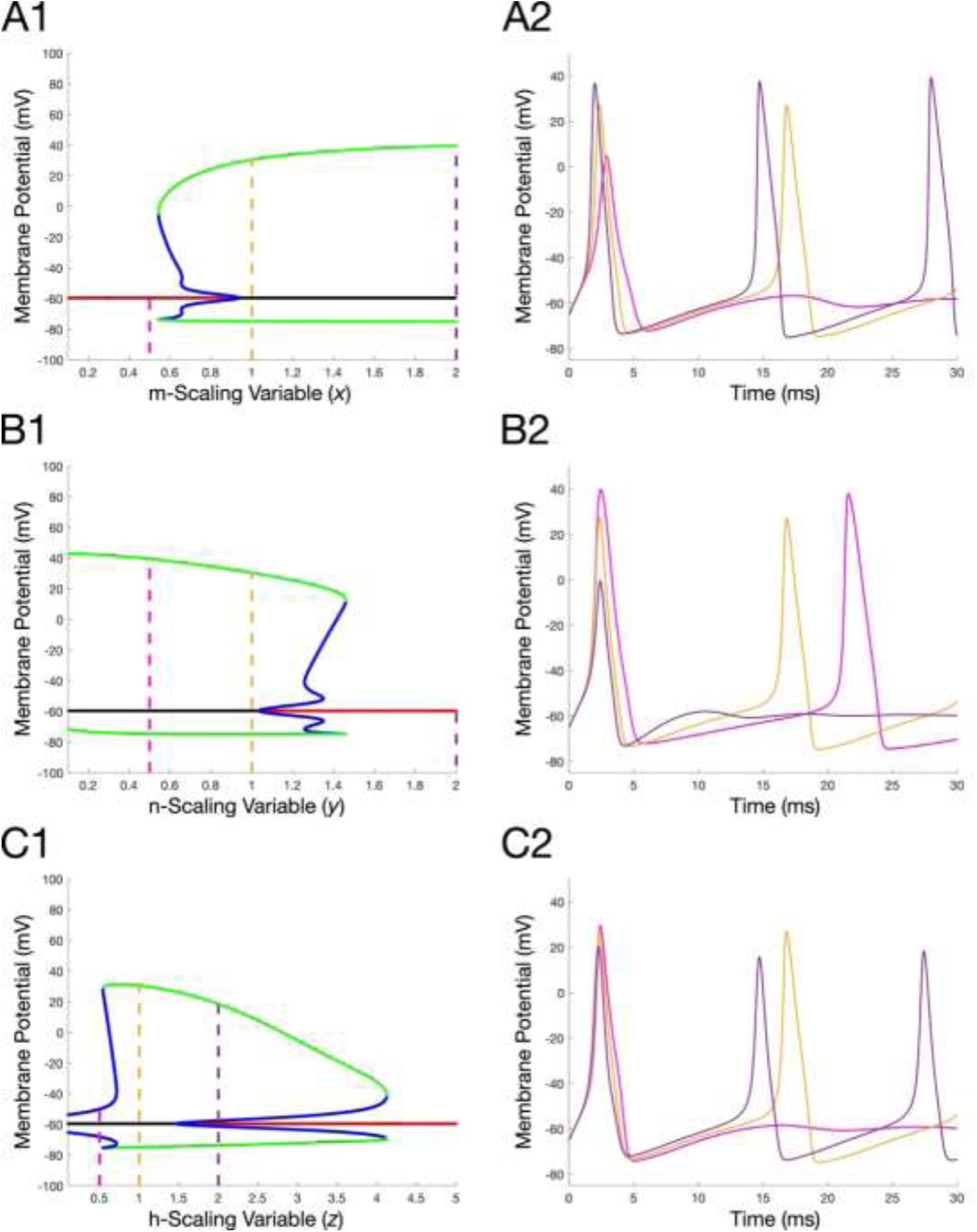
Bifurcation diagrams of the three scaling factors 𝑥 (**A1**), 𝑦 (**B1**), and 𝑧 (**C1**). The membrane potential trajectories are depicted for the values 0.5 (pink), 1 (yellow), and 2 (purple) for each of the parameters, x (**A2**), y (**B2**) and z (**C2**), illustrated by the dashed lines on the bifurcation diagrams. Note that all trajectories begin from the same initial condition (𝑉, 𝑚, 𝑛, ℎ) = (−65,0.1,0.1,0.1). Here, all simulations were carried by setting 𝐼_𝑠𝑡𝑖𝑚_ to 10 𝑝𝐴.

Increasing the K^+^ activation variable scaling factor (𝑦, Fig. 7B1) has the opposite effect on action potential generation and model dynamics than that of the Na^+^ activation variable scaling factor (𝑥, Fig. 7A). For small values of 𝑦 (for example, 0.5 in pink, or even 1 in yellow), the system is periodic and monostable with a stable limit cycle (surrounding an unstable equilibrium point), where model neurons generate sustained firing. As y is increased, the amplitude of the action potentials decreases as illustrated by the decrease in the upper periodic branch (as well as in Fig. 7B2, as described next). The equilibrium point eventually gains its stability via a subcritical Hopf bifurcation at 𝑦 = 1.038. The unstable limit cycle branch bifurcates outwards than coalesces with the stable limit cycle branch at an LPC point at 𝑦 = 1.46, reverting the system to monostability with a stable equilibrium point. Again, this allows for a transient interval of bistability 1.038 < 𝑦 < 1.46 in which two stable attractors coexist. These dynamical shifts are further justified in Fig. 7B2: If 𝑛 evolves more quickly and the repolarizing potassium current is established sooner, repolarization occurs before the membrane potential can reach 𝐸_𝑁𝑎_, lowering peak potential values. This is clearly illustrated by comparing the peak action potentials of the membrane potential trajectories elicited for 𝑦 = 0.5 (pink), 𝑦 = 1 (yellow), and 𝑦 = 2 (purple) curves. Similarly, if 𝑛 evolves more slowly, the depolarizing sodium current dominates the neuron’s dynamics for much longer, increasing peak values of action potentials. It is important to note, however, that increasing 𝑦 does shorten the repolarization process and thus increases the frequency of firing, as could be seen by comparing the 𝑦 = 0.5 and 𝑦 = 1 traces (Fig. 7B2).

Finally, Fig. 7C1 shows the bifurcation diagram for the Na^+^ inactivation variable scaling factor 𝑧. At low 𝑧 values (0.5413 < 𝑧 < 1.472), the system is monostable, with a stable limit cycle acting as the unique attractor and the model neuron is firing in a sustained mode. The interval of monostability ends via a subcritical Hopf bifurcation, which stabilizes the equilibrium at 𝑧 = 1.472 and creates a branch of unstable limit cycles. The unstable limit cycle branch bifurcates outwards and terminates the stable limit cycle at a LPC bifurcation at 𝑧 = 4.126. Similar to the effects of increasing 𝑦 on action potentials, increasing 𝑧 causes a decrease in the amplitude as well as the peak of the action potentials. Fig. 7C2 shows two trends illustrating the effect of 𝑧 on action potential morphology. First, as 𝑧 increases, the faster kinetics of ℎ causes it to recover back to its steady state value near 1 faster during hyperpolarization, shortening the refractory period and thus increasing the firing frequency. In addition, as 𝑧 increases the neuron becomes less excitable as faster sodium current inhibition impedes the depolarization mechanism. This is reflected in the reduction of the amplitude of the stable limit cycle as 𝑧 increases, followed by the disappearance entirely of the stable limit cycle branch at the 𝑧 = 4.126 LPC point.

Figure 8 expands our analysis to codimension-2 phenomena, here shown for the Na^+^ inactivation variable scaling factor (𝑧) and for the K^+^ activation variable scaling factor (𝑦) as parameters, while fixing the Na^+^ activation variable scaling factor (𝑥) to three different values (𝑥 = 1 in blue, 𝑥 = 2 in yellow, and 𝑥 = 4 in red). The effect of 𝑦 on the bifurcation diagrams of 𝑧 can be summarized by the fact that increased 𝑦 values impede action potentials by accelerating the onset of the depolarizing potassium current. This is reflected by the gradual contraction in the limit cycle region, which begins at 0.2185 < 𝑧 < 4.022 when 𝑦 = 0.5, contracts to 0.5415 < 𝑧 < 4.126 when 𝑦 = 1, and disappears entirely when 𝑦 = 2. In other words, at 𝑦 values near 2 and above, the loss of excitability due to accelerated potassium gating kinetics cannot be compensated by changing the dynamics of sodium inhibition. At high values of 𝑦, the primary cause of the loss of excitability is due to the fact that potassium-induced repolarization occurs immediately after the onset of the depolarizing sodium current, providing too small of a window to allow the neuron to sufficiently depolarize. As ℎ plays no role in this interplay of sodium and potassium activation, changing 𝑧 cannot correct this dynamical distortion.

The insets of Fig. 8 show the bifurcation diagrams for three different values of y (0.5, 1 and 2), while fixing 𝑥 to 1 (blue curve in the 2-parameter bifurcation diagram). Hopf bifurcations are illustrated via circles on the 2-param bifurcation diagram as well as on the 1-param diagrams in the insets. There is one generalized Hopf bifurcation point at which the first Lyapunov coefficient vanishes (shown as * on the diagram); we can assume the curve of LPC bifurcations terminates at this GH point. As described, increasing the value of 𝑦 causes a gradual contraction in the limit cycle region by accelerating potassium-induced repolarization, as evidenced by the reduction in the 𝑧 value at which the subcritical Hopf bifurcation occurs. Moreover, increasing 𝑥 accelerates the activation of depolarizing sodium currents and makes the neuron more excitable, such that the subcritical Hopf bifurcation along the 𝑦 -axis (near which the limit cycle region terminates) can be seen occurring at much higher 𝑦 values. Generally speaking, the region of the 𝑦 − 𝑧 plane at which the stable limit cycle is the unique system attractor (the region enclosed by the Hopf curve) expands as 𝑥 increases due to higher neuron excitability.

**Figure 8.**
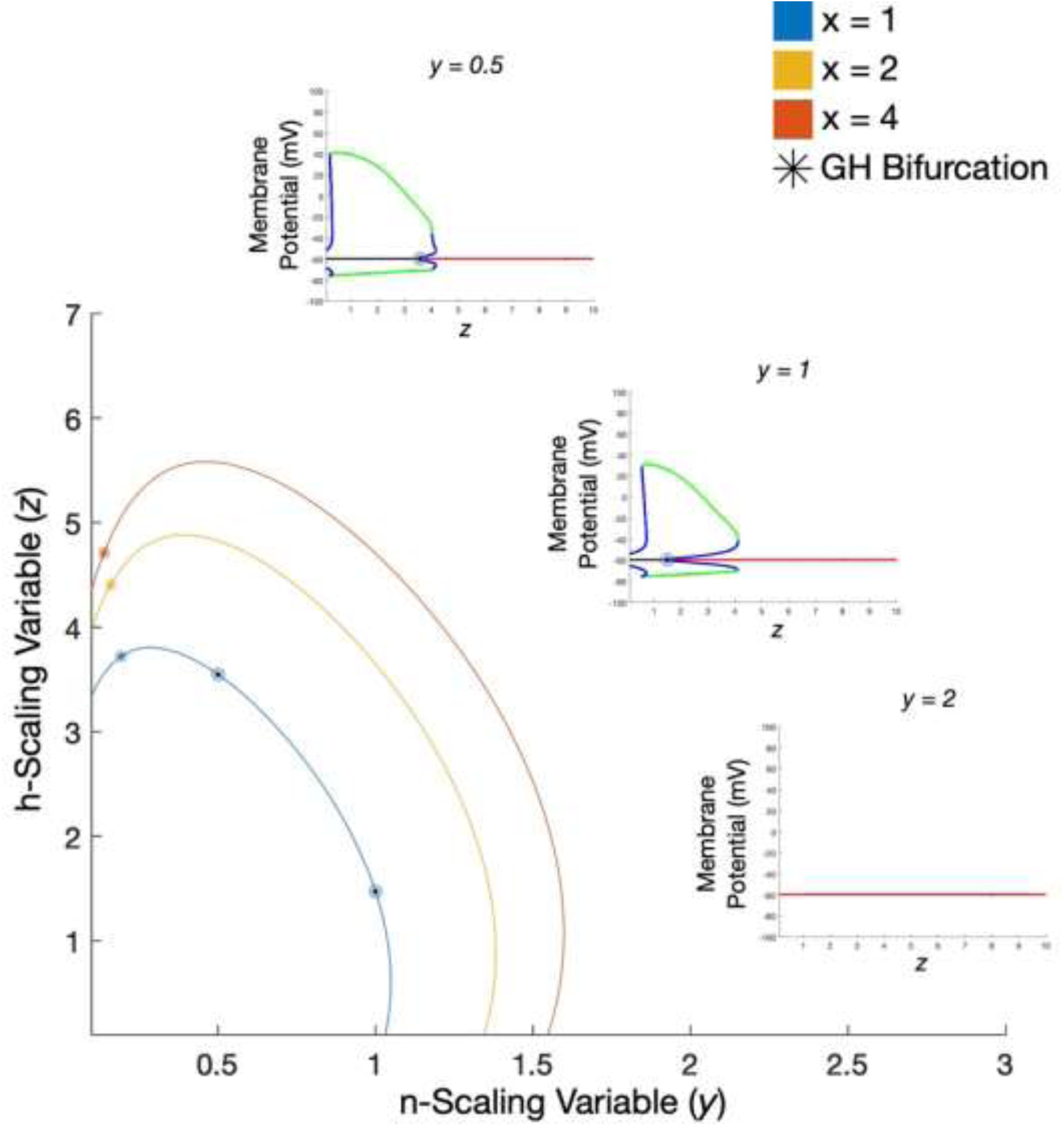
Overlay of codimension-2 Hopf point curves of 𝑦 and 𝑧 at various 𝑥 values (1 in blue, 2 in yellow, and 4 in red). Each of the insets correspond to a bifurcation diagram of 𝑧 at the fixed value 𝑥 = 1 and various 𝑦 values: 𝑦 = 0.5 (top left), 𝑦 = 1 (middle), and 𝑦 = 2 (bottom right). Circles illustrate Hopf bifurcation points on the co-dimension 2 curve (for 𝑥 = 1) and on the co-dimension 1 bifurcation diagrams. Generalized Hopf (GH) bifurcations are depicted with stars (*).

## Discussion

This study provides a comprehensive dynamical systems analysis of the temperature sensitivity of the classical Hodgkin-Huxley (HH) neuronal model, offering new mechanistic insights into how temperature perturbations alter neuronal excitability, spike morphology, and bifurcation structure. The bifurcations analysis was performed on an augmented Hodgkin-Huxley model that incorporated temperature via Arrhenius-like exponential relations between the transition rate constants of channel gating variables and temperature [19]. By integrating experimentally derived Q10 values into the HH framework and systematically varying temperature in conjunction with key membrane parameters, we have mapped the codimension-1 and codimension-2 bifurcation phase plots, identified generalized Hopf (Bautin) points, and generalized these findings to a broader class of temperature-dependent timescale modifications. We synthesized one and two-parameter bifurcation plots, trajectories on phase planes, and plots of individual morphological spike properties (spike amplitude, threshold potential, etc.) against temperature to examine how temperature, in conjunction with other standard HH model parameters, modulates the firing patterns of individual neurons. Additional analysis was carried out by incorporating separate timescale-distorting parameters (x, y, and z) for each ion channel to better isolate and identify the cause of shifts in spike morphology.

Our results confirm and extend previous experimental observations that temperature exerts dual, and sometimes opposing, effects on neuronal firing: it can accelerate spike generation via faster gating kinetics, but it can also suppress firing altogether once the balance between depolarizing and repolarizing currents is disrupted. We showed that temperature effects on the sodium activation variable *m* dominate at lower temperatures. The faster establishment of the depolarizing sodium current leads to substantially steeper depolarizations as well as higher action potential amplitudes due to a longer depolarization window. Beyond temperatures of 40 to 50 degrees; however, the acceleration of the kinetics of *n* and *h* become more prominent. Although firing frequency continues to increase due to the faster evolution of gating variables, the amplitude of action potentials substantially decrease as the earlier onset of the repolarizing potassium current and faster sodium deactivation provides a smaller window for the neuron to depolarize. At sufficiently high temperatures, these effects eliminate the stable limit cycle of the system entirely, making it impossible to sustain action potential chains due to the overpowering influence of faster *n* and *h* dynamics.

At low to moderate temperatures, accelerated 𝑚, ℎ, and 𝑛 kinetics shorten action potential width and refractory period, increasing firing frequency at the expense of spike amplitude. At higher temperatures, the breakdown of timescale separation between sodium and potassium gating eliminates the conditions for sustained oscillations, shown by the disappearance of stable limit cycles beyond a critical temperature. This “temperature-induced depolarization block” mirrors physiological findings in invertebrates and vertebrates, where excessive heating impairs axonal conduction and disrupts information transmission [5,44,45]. Importantly, we show that while temperature affects the various dynamical regimes explored, it leaves the position of equilibrium points invariant, highlighting that the primary effect is on the stability of solutions and not their fixed-point locations. This invariance is due to the fact that temperature rescales gating rates multiplicatively without altering the steady-state voltage-dependent relationships.

Bifurcation analyses revealed that increasing temperature contracts the range of parameters supporting sustained oscillations, both in terms of applied current and intrinsic conductances. In particular, for every parameter shown in Figure 6, increases in temperature beyond a certain level led to a contraction in the limit cycle region until the system becomes entirely monostable after the termination of the Hopf curve at a codimension-2 generalized Hopf point. For applied current in particular, the rightward movement of the subcritical Hopf bifurcation along the Istim axis is indicative of the neuron’s lower excitability at high temperatures due to faster *n* and *h* kinetics. The leftward movement of the supercritical Hopf point, meanwhile, suggests these augmented dynamics exacerbate the depolarization block phenomenon. Notably, higher temperatures increased the curvature of the unstable limit cycle curve, creating a larger region of bistability in which neurons are highly signal-sensitive as initial conditions can drastically change long-term behavior. Physiologically, this contraction implies a reduced operational range for neurons to maintain repetitive firing under thermal stress. Moreover, the parameter zones where both a stable equilibrium and a stable limit cycle coexist, renders the system more sensitive to small perturbations. In a biological context, such bistability could underlie abrupt state transitions in neural circuits during temperature fluctuations, potentially explaining observed shifts between tonic and silent states in sensory or central pattern generator networks.

By mapping temperature against standard HH parameters, we identified generalized Hopf bifurcations as organizing centers for the disappearance of oscillations. Above the GH point, the system is strictly monostable, regardless of applied current or conductance values. This provides a clear theoretical boundary for temperature tolerance in excitable membranes, a boundary that may correspond to experimentally observed thermal limits in neural tissues. These results largely align with previous literature on how temperature changes the action potential mechanisms of neurons. Ma et al. (2023), for instance, carried out *in vitro* experiments that showed increased firing frequencies and more sustainable action potential trains when temperatures were raised from 10 to 30 degrees in GFP-positive type II taste bud cells [46], corresponding to the limit cycle region of Figure 3 in which firing frequency increases with moderate decreases in spike amplitude. Simulations by Yu et al. (2012) that focused on temperature effects on the metabolic efficiency of axons found that increases in temperature reduced the duration of spike afterhyperpolarizations and spike amplitude [47]. This can be justified by faster *h* kinetics reducing neurons’ refractory periods and, with faster sodium inactivation, inhibiting depolarization. That being said, although our analysis provides a satisfying depiction of single-neuron systems, further study is needed to translate these results to neural circuits and systems.

Our results have several implications for both physiology and computational modeling. From a biological perspective, they suggest that species-specific adaptations to temperature, through channel isoforms with different Q10 values or altered gating kinetics, may be critical for maintaining neural performance in variable environments. From a modeling perspective, the detailed bifurcation maps and scaling-factor analysis provide a blueprint for extending the HH model to more realistic, heterogeneous neural populations and for exploring how temperature interacts with other modulatory factors such as pH, oxygenation, or pharmacological agents.

## References

1. Burrows M. Effects of temperature on a central synapse between identified motor neurons in the locust. J Comp Physiol. 1989;165: 687–695. doi:10.1007/BF00611000

2. Dalton JC, Hendrix DE. Effects of temperature on membrane potentials of lobster giant axon. American Journal of Physiology-Legacy Content. 1962;202: 491–494. doi:10.1152/ajplegacy.1962.202.3.491

3. Frankenhaeuser B, Moore LE. The effect of temperature on the sodium and potassium permeability changes in myelinated nerve fibres of Xenopus laevis. J Physiol. 1963;169: 431–437. doi:10.1113/jphysiol.1963.sp007269

4. Heitler WJ, Edwards DH. Effect of Temperature on a Voltage-Sensitive Electrical Synapse in Crayfish. Journal of Experimental Biology. 1998;201: 503–513. doi:10.1242/jeb.201.4.503

5. Hodgkin AL, Katz B. The effect of temperature on the electrical activity of the giant axon of the squid. The Journal of Physiology. 1949;109: 240–249. doi:10.1113/jphysiol.1949.sp004388

6. Kerkut GA, Ridge RMAP. The effect of temperature changes on the activity on the neurones of the snail Helix aspersa. Comparative Biochemistry and Physiology. 1962;5: 283–295. doi:10.1016/0010-406X(62)90057-9

7. Sjodin RA, Mullins LJ. Oscillatory behavior of the squid axon membrane potential. The Journal of General Physiology. 1958;42: 39–47. doi:10.1085/jgp.42.1.39

8. Gorman ALF, Marmor MF. Contributions of the sodium pump and ionic gradients to the membrane potential of a molluscan neurone. The Journal of Physiology. 1970;210: 897–917. doi:10.1113/jphysiol.1970.sp009248

9. Westerfield M, Joyner RW, Moore JW. Temperature-sensitive conduction failure at axon branch points. Journal of Neurophysiology. 1978;41: 1–8. doi:10.1152/jn.1978.41.1.1

10. Schoepfle GM, Erlanger J. The action of temperature on the excitability, spike height and configuration, and the refractory period observed in the responses of single medullated nerve fibers. American Journal of Physiology-Legacy Content. 1941;134: 694–704. doi:10.1152/ajplegacy.1941.134.4.694

11. Bittar P, Muller D. Time-dependent reversal of long-term potentiation by brief cooling shocks in rat hippocampal slices. Brain Research. 1993;620: 181–188. doi:10.1016/0006-8993(93)90154-F

12. Kim JA, Connors BW. High temperatures alter physiological properties of pyramidal cells and inhibitory interneurons in hippocampus. Front Cell Neurosci. 2012;6. doi:10.3389/fncel.2012.00027

13. Schauf CL. Temperature dependence of the ionic current kinetics of *Myxicola* giant axons. The Journal of Physiology. 1973;235: 197–205. doi:10.1113/jphysiol.1973.sp010384

14. Schwarz JR, Eikhof G. Na currents and action potentials in rat myelinated nerve fibres at 20 and 37° C. Pflugers Arch - Eur J Physiol. 1987;409: 569–577. doi:10.1007/BF00584655

15. Huxley AF. Ion movements during nerve activity. Annals of the New York Academy of Sciences. 1959;81: 221–246. doi:10.1111/j.1749-6632.1959.tb49311.x

16. Janssen R. Thermal influences on nervous system function. Neuroscience & Biobehavioral Reviews. 1992;16: 399–413. doi:10.1016/S0149-7634(05)80209-X

17. Moran O, Melani R. Temperature-dependent conduction properties in Arctic fish peripheral nerves. Polar Biology. 2001;24: 9–15. doi:10.1007/s003000000166

18. Swadlow HA, Waxman SG, Weyand TG. Effects of variations in temperature on impulse conduction along nonmyelinated axons in the mammalian brain. Experimental Neurology. 1981;71: 383–389. doi:10.1016/0014-4886(81)90096-0

19. Hodgkin AL, Huxley AF, Katz B. Measurement of current-voltage relations in the membrane of the giant axon of Loligo. The Journal of Physiology. 1952;116: 424–448. doi:10.1113/jphysiol.1952.sp004716

20. Sterratt DC. Q10: the Effect of Temperature on Ion Channel Kinetics. In: Jaeger D, Jung R, editors. Encyclopedia of Computational Neuroscience. New York, NY: Springer New York; 2015. pp. 2551–2552. doi:10.1007/978-1-4614-6675-8_236

21. Andersen P, Moser EI. Brain temperature and hippocampal function. Hippocampus. 1995;5: 491–498. doi:10.1002/hipo.450050602

22. Moser E, Mathiesen L, Andersen P. Association Between Brain Temperature and Dentate Field Potentials in Exploring and Swimming Rats. Science. 1993;259: 1324–1326. doi:10.1126/science.8446900

23. Fujii S, Sasaki H, Ito K, Kaneko K, Kato H. Temperature Dependence of Synaptic Responses in Guinea Pig Hippocampal CA1 Neurons in Vitro. Cell Mol Neurobiol. 2002;22: 379–391. doi:10.1023/A:1021068919709

24. Schiff SJ, Somjen GG. The effects of temperature on synaptic transmission in hippocampal tissue slices. Brain Research. 1985;345: 279–284. doi:10.1016/0006-8993(85)91004-2

25. Wu J, Javedan SP, Ellsworth K, Smith K, Fisher RS. Gamma oscillation underlies hyperthermia-induced epileptiform-like spikes in immature rat hippocampal slices. BMC Neurosci. 2001;2: 18. doi:10.1186/1471-2202-2-18

26. Wu J, Fisher RS. Hyperthermic Spreading Depressions in the Immature Rat Hippocampal Slice. Journal of Neurophysiology. 2000;84: 1355–1360. doi:10.1152/jn.2000.84.3.1355

27. Thompson S, Masukawa L, Prince D. Temperature dependence of intrinsic membrane properties and synaptic potentials in hippocampal CA1 neurons in vitro. J Neurosci. 1985;5: 817–824. doi:10.1523/JNEUROSCI.05-03-00817.1985

28. Alonso LM, Marder E. Temperature compensation in a small rhythmic circuit. eLife. 2020;9: e55470. doi:10.7554/eLife.55470

29. Long MA, Fee MS. Using temperature to analyse temporal dynamics in the songbird motor pathway. Nature. 2008;456: 189–194. doi:10.1038/nature07448

30. Burek M, Follmann R, Rosa E. Temperature effects on neuronal firing rates and tonic-to-bursting transitions. Biosystems. 2019;180: 1–6. doi:10.1016/j.biosystems.2019.03.003

31. Aronov D, Veit L, Goldberg JH, Fee MS. Two Distinct Modes of Forebrain Circuit Dynamics Underlie Temporal Patterning in the Vocalizations of Young Songbirds. J Neurosci. 2011;31: 16353–16368. doi:10.1523/JNEUROSCI.3009-11.2011

32. Aronov D, Fee MS. Analyzing the dynamics of brain circuits with temperature: Design and implementation of a miniature thermoelectric device. Journal of Neuroscience Methods. 2011;197: 32–47. doi:10.1016/j.jneumeth.2011.01.024

33. Banerjee A, Egger R, Long MA. Using focal cooling to link neural dynamics and behavior. Neuron. 2021;109: 2508–2518. doi:10.1016/j.neuron.2021.05.029

34. Goldin MA, Alonso LM, Alliende JA, Goller F, Mindlin GB. Temperature Induced Syllable Breaking Unveils Nonlinearly Interacting Timescales in Birdsong Motor Pathway. Coleman MJ, editor. PLoS ONE. 2013;8: e67814. doi:10.1371/journal.pone.0067814

35. Hamaguchi K, Tanaka M, Mooney R. A Distributed Recurrent Network Contributes to Temporally Precise Vocalizations. Neuron. 2016;91: 680–693. doi:10.1016/j.neuron.2016.06.019

36. Long MA, Katlowitz KA, Svirsky MA, Clary RC, Byun TM, Majaj N, et al. Functional Segregation of Cortical Regions Underlying Speech Timing and Articulation. Neuron. 2016;89: 1187–1193. doi:10.1016/j.neuron.2016.01.032

37. Okobi DE, Banerjee A, Matheson AMM, Phelps SM, Long MA. Motor cortical control of vocal interaction in neotropical singing mice. Science. 2019;363: 983–988. doi:10.1126/science.aau9480

38. Petersen PC, Vöröslakos M, Buzsáki G. Brain temperature affects quantitative features of hippocampal sharp wave ripples. Journal of Neurophysiology. 2022;127: 1417–1425. doi:10.1152/jn.00047.2022

39. Zhang YS, Wittenbach JD, Jin DZ, Kozhevnikov AA. Temperature Manipulation in Songbird Brain Implicates the Premotor Nucleus HVC in Birdsong Syntax. J Neurosci. 2017;37: 2600– 2611. doi:10.1523/JNEUROSCI.1827-16.2017

40. Hodgkin AL, Huxley AF. A quantitative description of membrane current and its application to conduction and excitation in nerve. The Journal of Physiology. 1952;117: 500–544. doi:10.1113/jphysiol.1952.sp004764

41. Dhooge A, Govaerts W, Kuznetsov YuA. MATCONT: a Matlab package for numerical bifurcation analysis of ODEs. SIGSAM Bull. 2004;38: 21–22. doi:10.1145/980175.980184

42. Strogatz SH. Nonlinear Dynamics and Chaos: With Applications to Physics, Biology, Chemistry, and Engineering. 3rd ed. Boca Raton: Chapman and Hall/CRC; 2024. doi:10.1201/9780429398490

43. Ermentrout B. Type I Membranes, Phase Resetting Curves, and Synchrony. Neural Computation. 1996;8: 979–1001. doi:10.1162/neco.1996.8.5.979

44. Austerschmidt LJ, Khan A, Plant DO, Richards EMB, Knott S, Baker MD. The effects of temperature on the biophysical properties of optic nerve F-fibres. Sci Rep. 2020;10: 12755. doi:10.1038/s41598-020-69728-y

45. Ding Q, Jia Y. Effects of temperature and ion channel blocks on propagation of action potential in myelinated axons. Chaos: An Interdisciplinary Journal of Nonlinear Science. 2021;31: 053102. doi:10.1063/5.0044874

46. Ma Z, Paudel U, Foskett JK. Effects of temperature on action potentials and ion conductances in type II taste-bud cells. Am J Physiol Cell Physiol. 2023;325: C155–C171. doi:10.1152/ajpcell.00413.2022

47. Yu Y, Hill AP, McCormick DA. Warm Body Temperature Facilitates Energy Efficient Cortical Action Potentials. Sporns O, editor. PLoS Comput Biol. 2012;8: e1002456. doi:10.1371/journal.pcbi.1002456

